# Evolution of anticipatory effects mediated by epigenetic changes

**DOI:** 10.1101/2020.09.30.321091

**Authors:** Ilkka Kronholm

## Abstract

Anticipatory effects mediated by epigenetic changes occur when parents modify the phenotype of their offspring by making epigenetic changes in their gametes guided by information from an environmental cue. To investigate when do anticipatory effects mediated by epigenetic changes evolve in a fluctuating environment, I use an individual based simulation model with explicit genetic architecture. The model allows for the population to respond to environmental changes by evolving plasticity, bet-hedging, or by tracking the environment with genetic adaptation, in addition to the evolution of anticipatory effects. The results show that anticipatory effects evolve when the environmental cue provides reliable information about the environment and the environment changes at intermediate rates, provided that fitness costs of anticipatory effects are rather low. Moreover, evolution of anticipatory effects is quite robust to different genetic architectures when reliability of the environmental cue is high. Anticipatory effects always give smaller fitness benefits than within generation plasticity, suggesting a possible reason for generally small observed anticipatory effects in empirical studies.

## Introduction

Some organisms are able to prime their offspring for future environmental challenges that the offspring are likely to face. These effects have been called anticipatory parental effects, between generation plasticity, induced epigenetic changes, or transgenerational effects in the literature (Jablonka & Raz, 2009; Marshall & Uller, 2007; Kronholm, 2017). From now on, I will use the term anticipatory effects following Marshall & Uller (2007), to focus on the phenotypic consequences and to remain agnostic about the molecular mechanism responsible for the effect. The clearest examples of anticipatory effects have been observed in plants, if parents have encountered pathogens or herbivores they prime their offspring to have elevated levels of defense against pathogens (Rasmann et al., 2012; Slaughter et al., 2012; Luna et al., 2012) or herbivores (Holeski, 2007). Offspring can also be prepared to deal with abiotic factors such as drought and osmotic stress (Wibowo et al., 2016; Herman et al., 2012) or shading (Galloway & Etterson, 2007). Anticipatory effects are not restricted to plants, but have been observed in many animals as well. Particularly, there are examples of fish responding to temperature during development in subsequent generations (Salinas & Munch, 2012; Donelson et al., 2012; Shama & Wegner, 2014), water fleas developing different morphs in response to predators (Agrawal et al., 1999), nematodes respond to viral infection and starvation in subsequent generations (Rechavi et al., 2011, 2014; Jobson et al., 2015; Kishimoto et al., 2017; Ivimey-Cook et al., 2020), and metabolic changes in offspring in response to diet in fruit flies (Öst et al., 2014). A general principle in these examples is that the parents experience a particular environment, and as a result they change the phenotype of their offspring in a presumably adaptive manner. Priming the offspring to better cope with this particular environmental challenge, even if the offspring have not yet encountered this environment themselves.

Anticipatory effects require a mechanism that makes it possible, for example, for offspring to upregulate the expression of possibly costly defence related genes, even if they have not yet encountered a particular pathogen. There are different mechanisms that may accomplish this, such as provisioning of nutrients or hormones to eggs, transfer of antimicrobial peptides or antibodies to the offspring, or epigenetic inheritance. In some cases the mechanistic basis of anticipatory effects based on epigenetic inheritance has been worked out in great detail. The specifics vary by taxon: in the plant *Arabidopsis thaliana* certain genes are methylated by the RNA-dependent methylation pathway as a response to osmotic stress, and these methylation patterns can be inherited via seed but not by pollen (Wibowo et al., 2016). In the nematode *Caenorhabditis elegans* environmental stress induces the production of small RNA’s that are inherited, and these RNA’s subsequently direct chromatin conformation and gene expression changes of their target genes (Rechavi et al., 2014; Rechavi & Lev, 2017). In fruit flies epigenetic inheritance was dependent on the Polycomb repressive complex silencing machinery (Öst et al., 2014), and small RNA’s may be involved in fruit flies as well (Duempelmann et al., 2020).

In ordinary phenotypic plasticity organisms use some sort of environmental cue to inform delevopmental decisions. For instance, changes in day length predict the change of seasons and thus temperature. Conceptually, anticipatory effects closely resemble ordinary phenotypic plasticity that happens within a generation, but the inducing environmental signal has been perceived by the parent rather than the focal organism. In order for anticipatory effects to occur, there needs to be a genetic pathway already present that can sense, transduce, and respond to an environmental signal, like in normal within generation phenotypic plasticity. Intuitively, the evolution of the capability to use anticipatory effects seems to be contingent on the environmental cue having some predictive power on what the environment will be like in the next generation.

Several previous theoretical studies have investigated the evolution of maternal effects that are transmitted using models based on phenotypic memory, where the phenotype of the parent is partially passed on the to the offspring (Jablonka et al., 1995; Lachmann & Jablonka, 1996; Hoyle & Ezard, 2012; Prizak et al., 2014; Kuijper & Hoyle, 2015; Kuijper & Johnstone, 2015; Leimar & McNamara, 2015). Because genes influence the phenotype of the parent and the phenotype is passed partially to offspring, phenotypic memory can be considered a special case of indirect genetic effects, which can then cascade over multiple generations (Hoyle & Ezard, 2012; Kuijper & Hoyle, 2015). Previous studies indicate that the evolution of anticipatory effects is favoured in fluctuating environments (Jablonka et al., 1995; Lachmann & Jablonka, 1996; Kuijper & Johnstone, 2015; Leimar & McNamara, 2015). Degree of environmental autocorrelation and predictive accuracy of environmental cues influences whether maternal information is used (Leimar & McNamara, 2015). However, some models give different results as to what has to be the timescale of environmental fluctuations (Furrow & Feldman, 2014). Moreover, the evolution of anticipatory effects has been studied in the context of population structure, and moderate levels of gene flow were found to favour the evolution of anticipatory effects (Kuijper & Johnstone, 2015; Leimar & McNamara, 2015; Greenspoon & Spencer, 2018).

While the previous models have have shed light on the conditions where anticipatory effects should evolve, there are still some uncertanties. First, it has been suggested that organisms can use different types of strategies to adapt to environmental changes that happen at different time scales (Kristensen et al., 2020). Environmental changes that happen within a generation should favor the evolution of within generation plasticity, slightly longer fluctuations could favour anticipatory effects, and populations should adapt to environmental changes over long periods of time by genetic adaptation. Yet, the exact conditions that the favour the evolution of anticipatory effects in fluctuating environments when alternative strategies are also possible are unclear at the moment. There could be regions of the parameter space where anticipatory effects would evolve only if investigated in isolation but not when alternative strategies are also possible, as they might have a higher fitness preventing the evolution of anticipatory effects. Therefore, it is important to investigate the evolution of anticipatory effects in combination with alternative strategies, as done for phenotypic plasticity by Botero et al. (2015). Furthermore, a recent meta-analysis found that anticipatory effects are mostly weak (Uller et al., 2013). Whether it is the case that conditions for the evolution of anticipatory effects are restrictive such that they are rare in nature, or that those taxa that live in conditions that are favourable for the evolution of anticipatory effects are under represented in studies, or that anticipatory effects are generally to be expected to be low, is not known. By having a clear picture of the conditions where anticipatory effects are expected to evolve, empirical studies can focus on those species that live in such environments.

Second, previous models have mainly investigated the evolution of maternal or anticipatory effects using greatly simplified genetic architectures. Either based on a few loci with restricted allelic effects, or a quantitative genetic framework, where the assumption is that traits are determined by infinitely many small effect loci. While this is likely to be a reasonable assumption for many phenotypes, there are certainly examples of large genetic contributions of single loci to the phenotype for a number of different traits, and if genetic architecture is based on a few large effect loci, this can change the observed evolutionary dynamics (Oomen et al., 2020). There could be constraints on the evolution of anticipatory effects if, for example, mutational effects sizes or the number of loci are important factors for the evolution of the genetic traits that govern the induction of anticipatory effects.

Furthermore, while it is certainly possible epigenetic changes and indirect genetic effects based on phenotypic memory will conceptually work in a similar manner, this is not completely clear. In the case of epigenetic inheritance, parents can pass on information about their current environment in the form of chemical modifcations of DNA or associated histone proteins, and this information about the current environment can be different than the current phenotype of the parents.

I ask the question: in what kind of fluctuating environments do anticipatory effects evolve? I mainly focus on investigating how does the predictability of the environment and the frequency of environmental fluctuations affect the evolution of anticipatory effects and what is the effect of relaxing the assumption of infinitely many loci of small effects. Furthermore, I investigate what effect do costs of phenotypic plasticity and anticipatory effects have on their evolution, and does genetic architecture constrain the evolution of anticipatory effects? To answer these questions I use an individual based simulation model adapted and modified from Botero et al. (2015). In the model, each individual has a linear reaction norm that is determined by an explicit genetic architecture with arbitrary mutational effects. The model allows the population to evolve either reversible phenotypic plasticity where phenotype can be adjusted multiple times per generation, developmental plasticity where phenotype is adjusted only once, increased developmental variation as a type of bet-hedging strategy, or simple genetic adaptation to the environment by tracking the environmental optimum by changing reaction norm intercept. Finally, the population can also evolve epigenetic modification, allowing anticipatory effects, where information from the parents is passed on the the offspring that can adjust their phenotype in early development.

## Methods

### Simulation model

I used an individual based Wright-Fisher model with non-overlapping generations and random mating. The model keeps track of explicit genetics of each individual. Phenotype is determined by multiple loci and environmental effects. Environment changes at different rates and individuals can potentially use an environmental cue to respond by plasticity or by anticipatory parental effects mediated by epigenetic changes. Developmental variation is also partially under genetic control, which makes it possible to evolve increased variation as a bet-hedging type strategy. The model is based on a model of phenotypic plasticity used by Botero et al. (2015) with some modifications. The events in the model are shown in Figure 1.

**Figure 1:**
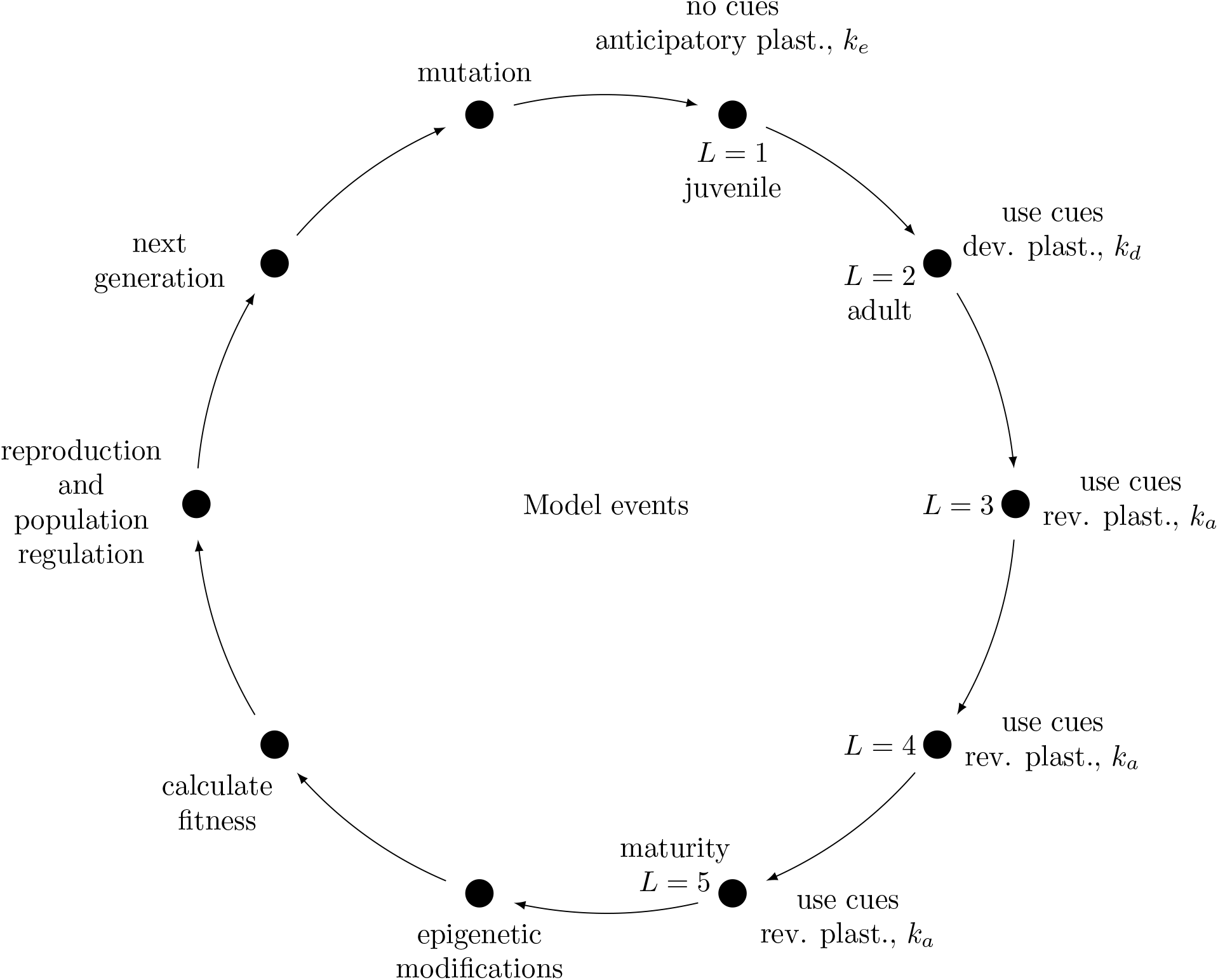
Events in the simulation model. Each individual lives for 5 timesteps. In the first timestep individuals can use epigenetic changes inherited from their parents to modify their phenotype and pay a fitness cost of *k*_*e*_ for this action. Epigenetic modifications are reset at the end of this step. In the next timestep individuals can use developmental plasticity to adjust their phenotype based on an observed environmental cue, with a fitness cost of *k*_*d*_. In timesteps three to five, individuals can use reversible plasticity to further adjust their phenotype based on observed cues, with a fitness of cost of *k*_*a*_ for each adjustment. After timestep five, any potential epigenetic modifications are set based on an observed environmental cue from timestep five. Then fitness is calculated for each individual, individuals produce offspring in proportion to their fitness, population regulation happens, and next generation is formed. Then new mutations are produced in those individuals and the cycle starts again.

#### Environmental changes

In the simulation, there are *L* timesteps in each generation, amounting to a total of *t* = *n*_*g*_*L* timesteps, where *n*_*g*_ is the number of generations. The number of timesteps in each generation was set to 5. The environment, *E*, changes cyclically according to:

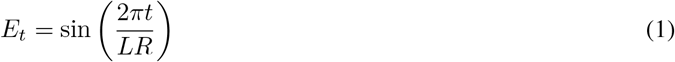

where *R* is a parameter that determines how fast the environment changes relative to generation time. Figure S1 shows how changing *R* affects the speed of environmental fluctuations. Small values of *R* cause the environment to fluctuate fast and large values of *R* mean slow fluctuations. The values of *R* that were used in the simulations were: 1, 3.2, 10, 31.6, 100, 316.2, 1000, 3162.3, and 10000. In subsequent plots these values are plotted as their base 10 logarithms: 0, 0.5, 1, 1.5, 2, 2.5, 3, 3.5, and 4.

There is also an environmental cue, *C*, that individuals can potentially use to anticipate changes in the environment. The distribution of the cue is gaussian: *C*_*t*_ ∼ N(*E*_*t*_*P*, (1 − *P*)*/*3), where *P* is a parameter that determines the predictability of the environment. When *P* = 1 the cue predicts the environment perfectly, and when *P* = 0, cue is randomly distributed around 0 with standard deviation of 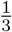. Simulations were run for values of *P* from 0 to 1 at 0.1 intervals. While the environment changes in a derterministic manner, the environment is not necessarely predictable from the point of view of the organism as the environmental cue that the organisms observe has randomness built in. Furthermore, the offspring do not necessarely experience the same environment as their parents did if the rate of environmental change is moderate relative to generation time. Botero et al. (2015) also tried other functions for *E* than a sine wave and observed that similar results were obtained, as the ability to predict the environment based on the cue is more important than the form of fluctuations that *E* has.

#### Genetics, development, and phenotype

Individuals’ phenotype, *Z*, is determined as in a standard quantitative genetic model, but with explicit loci. As the model allows for plasticity, the phenotype for individual *i* at time point *t* is determined by a linear reaction norm

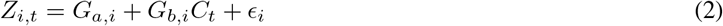

where *G*_*a*_ is the genotypic value for the intercept, *G*_*b*_ is the genotypic value for the slope by which an individual changes its phenotype as a reaction to the environmental cue, and ∈ which is random developmental variation in the phenotype. The random variation in development is distributed as ∈ ∼ N(0, *σ*_*R*_), where *σ*_*R*_ = *σ*_*E*_ + *G*_*h,i*_ and *σ*_*E*_ is the environmental component of random variation and *G*_*h*_ the genotypic component of random developmental variation. Thus, the amount of random developmental variation can potentially evolve. There are *l*_*a*_ loci that determine the intercept, *l*_*b*_ loci that determine the slope, *l*_*d*_ loci that determine the probability of plastic adjustment of the phenotype, *l*_*h*_ loci that determine the genotypic component of random developmental variation, and *l*_*e*_ loci that determine the probability of epigenetic modification of slope loci. Genotypic values are always the sum of individual allelic effects for each locus, for example 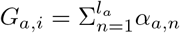. However, genotypic values for adjustment probability, *G*_*d*_, and probability of epigenetic modification, *G*_*e*_, were restricted between 0 and 1. The genotypic values for random developmental variation, *G*_*h*_, were restricted to between 0 and 3. Restricted genotypic values were adjusted by setting values of *G <* 0 to 0 and *G >* 1 to 1. The underlying allelic effects could evolve freely and were not restricted, this means that a population could evolve a robust probability of either 0 or 1, which seems biologically reasonable for probabilities. For *G*_*h*_ the limit was placed so that biologically completely unrealistic values could not evolve and I never observed values close to the upper limit in the simulations. There were no non-additive allelic effects. The number of loci for each category was set to 10 unless otherwise stated. At the start of the simulation, a genetic map was randomly generated, with all loci distributed randomly and uniformly across *n*_*c*_ linkage groups (chromosomes).

Individuals live for five timesteps (stages): during the first stage an individual has a chance to adjust its phenotype based on epigenetic modifications received from parents, but it cannot yet use environmental cues on its own. The anticipatory effect, 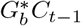 is calculated over slope loci if they are epigenetically modified, 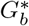 denotes an epigenetic slope effect. The phenotype of an individual that uses epigenetic information in its first life stage is 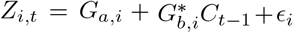. Modified alleles incorporate the allelic effect of slope and the cue experienced by the parents in timestep *t* − 1. In other words, modified alleles store in cue information experienced by the parents in the previous timestep. For an individual that has no epigenetic modifications the phenotype at first life stage is *Z*_*i,t*_ = *G*_*a,i*_ + *E*_*i*_. In the second life stage individuals can use developmental plasticity according to equation 2. In the third, fourth, and fifth life stage individuals can use reversible plasticity to adjust their phenotype based on environmental cues with a probability of *G*_*d*_ at each step, so individuals phenotype follows equation 2 if it adjusts, and *Z*_*i,t*_ = *G*_*a,i*_ + *E*_*i*_ if it does not. In the fifth and final life stage, individuals epigenetically modify all of their slope alleles with a probability of *G*_*e*_. For each slope allele there are two values stored: its allelic effect and epigenetic modification. If an individuals modifies its alleles epigenetically it modifies all of them setting the epigenetic modification to the value of *C*_*t*_. Then events of the model: reproduction and mutation happen as in figure 1. Then in the next generation at first life stage individuals have a chance to use the stored epigenetic information, which will be from time point *C*_*t−*1_, as explained above.

#### Fitness and selection

Fitness for each individual over its entire life span is calculated as the sum of phenotypic deviations from the environmental optimum at each life stage:

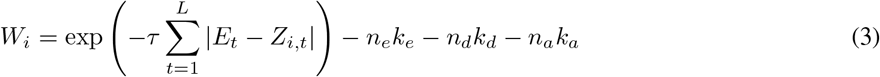

where *τ* is a parameter for intensity of selection which was set to 0.25, *k*_*e*_ is the fitness cost of adjusting the phenotype using epigenetic modifications, *n*_*e*_ ∈ {0, 1} is the number of such adjustments, *k*_*d*_ is the fitness cost for adjusting the phenotype by developmental plasticity, *n*_*d*_ ∈ {0, 1} is the number of such adjustments, *k*_*a*_ is the fitness cost for adjusting the phenotype by reversible phenotypic plasticity, and *n*_*a*_ ∈ {0, 1, 2, 3} is the number of such adjustments. Any negative values for fitness due to costs of phenotypic plasticity were set to 0.

#### Reproduction and population regulation

The number of offspring produced by individual *i* is drawn from a Poisson distribution with mean of *W*_*i*_*N*_*R*_, where *N*_*R*_ is the number of offspring produced on average by a perfectly adapted individual (that has a relative fitness of 1). Since individuals are hermaphrodites that cannot self, offspring are generated for individual *i* so that for each offspring the other parent is picked randomly with replacement from the population weighted by their fitness. Thus fitness affects how many offspring an individual produces and the probability to participate in matings. Gametes are produced by meiosis and the number of cross-overs for each pair of homologous chromosomes has a distribution of *n*_*xo*_ ∼ Pois(*λ*) + 1, so that the minimum number of cross-overs is always 1. The *λ* parameter was set to 0.56, which corresponds to 1.56 cross-overs for each pair of homologous chromosomes in meiosis. This is the number of chiasmata per chromosome pair that on average is observed across eukaryotes (Otto & Lenormand, 2002). Cross-over positions are drawn from a uniform distribution along the chromosome, with no interference. If and only if the total number of offspring produced by all individuals was larger than the population carrying capacity, *K*, then offspring were removed randomly until *K* offspring remained. Generally carrying capacity was reached in the simulations. Preliminary tests were also done with a different method of density regulation, where fitness affects the probability that individuals get to reproduce and then how many individuals they produce according to a logistic growth model. Method of density regulation had no significant effect on the results.

#### Mutation

After the next generation of individuals has been produced, mutations are generated. Each locus has a per locus mutation rate, *µ*. Since there are 2*Nl* alleles in a population, the mean number of mutations per generation is 2*Nlµ*. Each mutation generates a new allele and the allelic effect is drawn from a gaussian distribution, *α* ∼ N(0, *σ*_*α*_). Standard deviation of mutational effects, *σ*_*α*_, was set to 1. Mutational variance, which describes how much genetic variation is contributed by mutation in each generation, can be calculated as 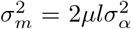, and mutational heritability by standardizing mutational variance with environmental variance, 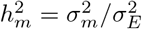 (Lynch & Walsh, 1998). Substituting values used in the simulation to these equations leads to a mutational heritability of 0.2 for the reaction norm intercept or slope. This value is one to two orders of magnitude higher than typical mutational heritabilities that have been measured experimentally (Halligan & Keightley, 2009; Durand et al., 2010; Lynch & Walsh, 1998), presumably because per locus mutation rate used in the simulation, *µ* = 10^*−*4^, is much higher than realistic mutation rates. However, these values of mutational parameters were used because of the relatively small population sizes that could be used in the simulation without extending the computational time beyond feasibility. Increased mutation rate may cause the simulations to reach an evolutionary optimum faster than would naturally happen, but this is not certain as population size in the simulations is much lower than in natural populations. Furthermore, mutation rates that are relevant for adaptation also depend on mutational target sizes, which are difficult to estimate. A large mutational target will cause the per locus mutation rate to be much higher than the per base pair rate. Robustness of the results to mutation rates will be examined below.

### Simulations and data analysis

Simulations were written in R (R Core Team, 2019). Replicate simulations were run in parallel on a computer super-cluster utilizing the ‘snow’ package. To examine in which conditions anticipatory effects evolve, I ran 100 replicate simulations for each combination of *R* and *P*. Costs of plasticity were set to *k*_*d*_ = *k*_*e*_ = 0.02 and *k*_*a*_ = 0.01, which correspond to values used by Botero et al. (2015) for the costs of developmental and reversible plasticity. Each replicate simulation lasted for 3000 generations. After these simulations it was clear that 3000 generations was more than enough for the populations to adapt, so in subsequent simulations 1500 generations were used to reduce computational time. The simulations were initialised so that there was no genetic variation present at the start. Each simulation started with 2000 individuals. Results were reported as the values of the different variables at the end of the simulation. All simulation parameters are shown in table 1. Then I tested how costs of plasticity affect the evolutionary outcomes. I ran simulations where there were no costs of phenotypic plasticity or anticipatory effects, *k*_*d*_ = *k*_*e*_ = *k*_*a*_ = 0 and simulations where the costs were twice as high as in the the original simulations *k*_*d*_ = *k*_*e*_ = 0.04 and *k*_*a*_ = 0.02.

**Table 1:**
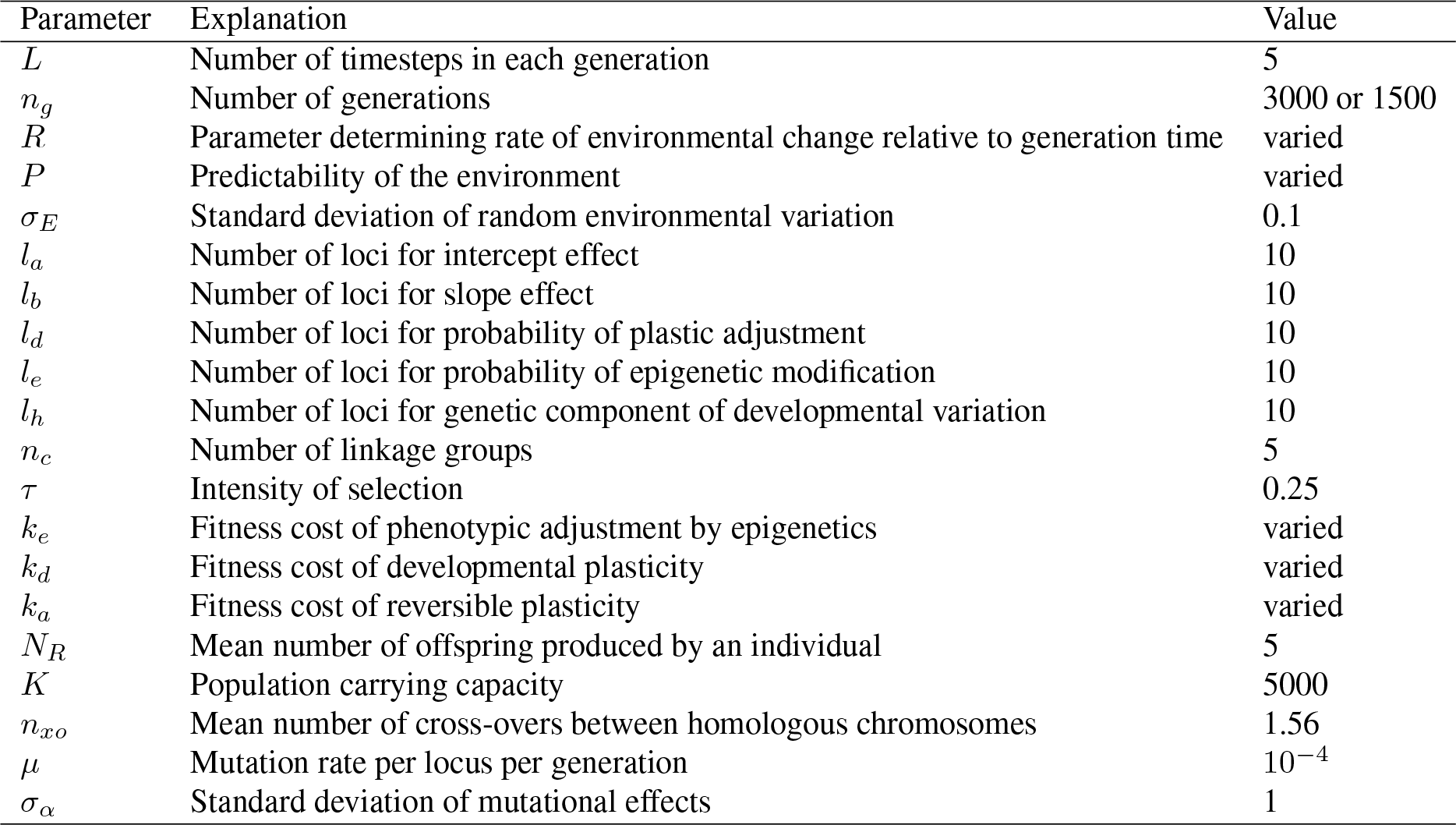
Parameters used in the simulation model.

| Parameter | Explanation | Value |
| --- | --- | --- |
| $L$ | Number of timesteps in each generation | 5 |
| $n_g$ | Number of generations | 3000 or 1500 |
| $R$ | Parameter determining rate of environmental change relative to generation time | varied |
| $P$ | Predictability of the environment | varied |
| $\sigma_E$ | Standard deviation of random environmental variation | 0.1 |
| $l_a$ | Number of loci for intercept effect | 10 |
| $l_b$ | Number of loci for slope effect | 10 |
| $l_d$ | Number of loci for probability of plastic adjustment | 10 |
| $l_e$ | Number of loci for probability of epigenetic modification | 10 |
| $l_h$ | Number of loci for genetic component of developmental variation | 10 |
| $n_c$ | Number of linkage groups | 5 |
| $\tau$ | Intensity of selection | 0.25 |
| $k_e$ | Fitness cost of phenotypic adjustment by epigenetics | varied |
| $k_d$ | Fitness cost of developmental plasticity | varied |
| $k_a$ | Fitness cost of reversible plasticity | varied |
| $N_R$ | Mean number of offspring produced by an individual | 5 |
| $K$ | Population carrying capacity | 5000 |
| $n_{xo}$ | Mean number of cross-overs between homologous chromosomes | 1.56 |
| $\mu$ | Mutation rate per locus per generation | $10^{-4}$ |
| $\sigma_\alpha$ | Standard deviation of mutational effects | 1 |

#### Sensitivity analysis

To examine if the results were robust to different genetic architectures, I ran a sensitivity analysis of the simulation. 100 replicate simulations for each parameter combination of *P* and *R* were run. For each replicate, random values were drawn for parameters *σ*_*E*_, number of different loci, linkage groups, mutation rate, and *σ*_*α*_. Costs of plasticity were *k*_*d*_ = *k*_*e*_ = 0.02 and *k*_*a*_ = 0.01. Other parameters were the same as in previous simulations (Table 1). Values were drawn from the following uniform distributions: *σ*_*E*_ ∼ U(0.01, 0.5), *µ* ∼ U(10^*−*6^, 10^*−*3^), *σ*_*α*_ ∼ U(0.01, 2), number of loci of was randomly selected from [4, 25] and was the same for each type of loci, and number of linkage groups from [1, 10]. Loci were distributed evenly across the linkage groups, and the remainder was always assigned to linkage group 1.

### Data availability

An R package implementing the simulations is available on github: https://github.com/ikron/indsim.

## Results

### Phenotypic plasticity

When costs of plasticity were *k*_*e*_ = *k*_*d*_ = 0.02 and *k*_*a*_ = 0.01, some sort of plasticity consistently evolved when the parameter determining rate of environmental change, *R*, was less than 100 (log_10_ *R <* 2) and environmental predictability, *P*, was 0.3 or greater (Figure 2). Reversible plasticity (Figure S2) evolves when rate of environmental change is fast, that is when R is 10 ((log_10_ *R* = 1) or smaller. Higher values of *R* favoured the evolution of developmental plasticity (Figure S3), thus when environmental change is slow enough the costs of reversible plasticity outweigh its fitness benefits. Developmental plasticity evolved in some replicates when *R* was 100 (log_10_ *R* = 2) and predictability high, but slower environmental change did no longer favour the evolution of plasticity. Instead the population tracked the moving environmental optimum by changing the reaction norm intercept (Figure S4). When environmental change was unpredictable (*P <* 0.3) and fast (*R* = 1, log_10_ *R* = 0), the reaction norm of the population remained flat with an intercept of 0 (Figure 2). This represents a conservative strategy where individuals are just tolerating environmental changes. However, when environmental changes were slower (*R* = 10, log_10_ *R* = 1) but still unpredictable, the population evolved increased developmental variation (Figure S5). This represents a bet-hedging strategy where individuals benefit from producing offspring with different phenotypes in order for some of them to be adaptated to the unpredictable environment. Overall the results regarding plasticity are very similar as those obtained by Botero et al. (2015), despite some differences in the genetic architecture of the traits. Here the conditions favouring the evolution of a diversifying bet-hedging strategy are more restrictive than in Botero et al. (2015), but in their model it was possible to evolve a phenotype that produced two distinct morphs, rather than only increased developmental variation as here.

**Figure 2:**
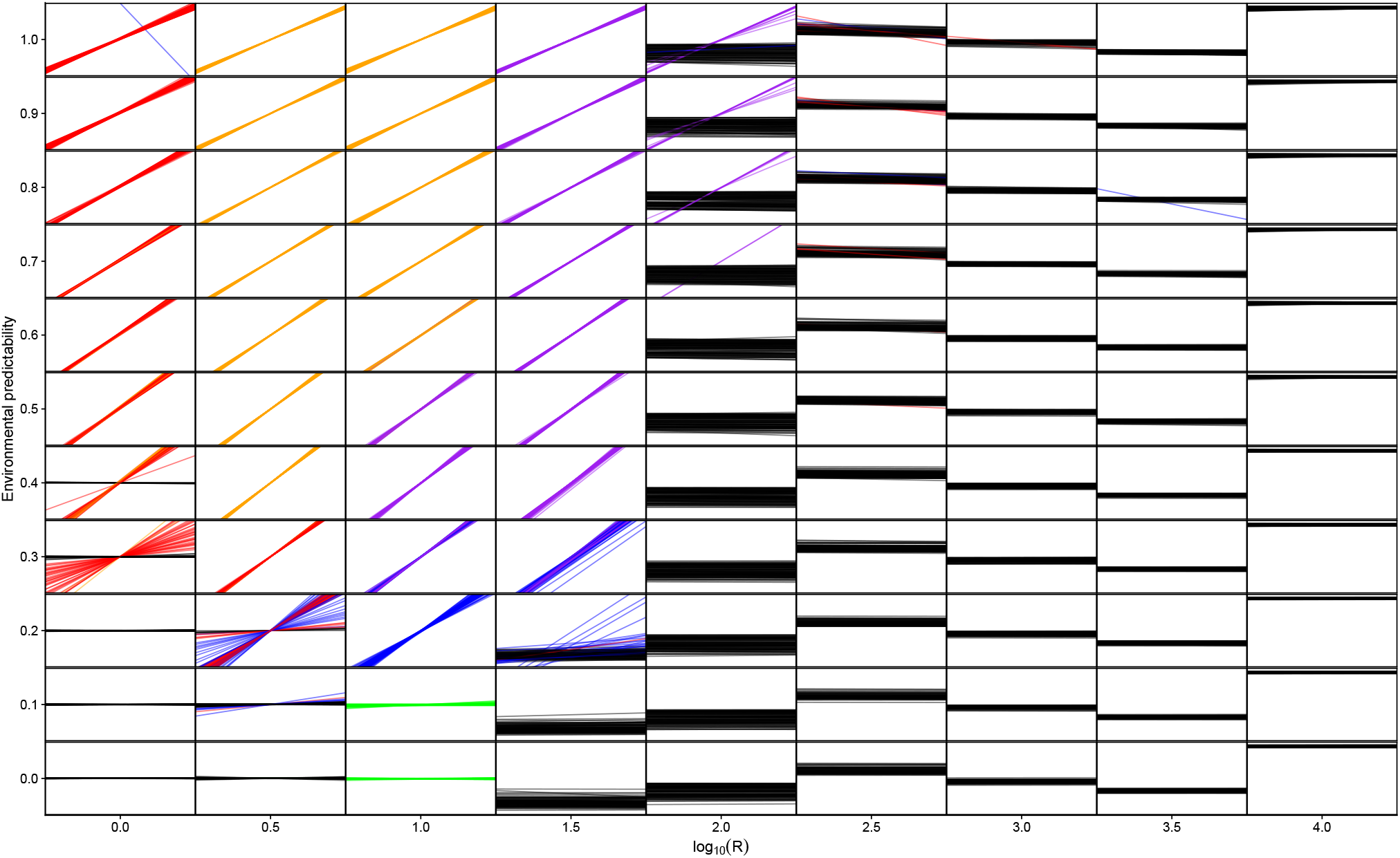
Simulation results for *k*_*e*_ = *k*_*d*_ = 0.02 and *k*_*a*_ = 0.01. For each small panel, the environmental cue is on the x-axis and phenotype on the y-axis. Each reaction norm is the population mean for one replicate simulation and 100 replicates were performed for each combination of *P* and *R*. Red colour shows reaction norms, where the population evolved reversible plasticity (*G*_*b*_ *>* −0.1 or *G*_*b*_ *<* −0.1, and *G*_*d*_ *>* 0.1). Orange colour shows reaction norms where reversible and anticipatory effects evolved (in addition *G*_*e*_ *>* 0.1), blue colour indicates that only developmental plasticity evolved (*G*_*b*_ *>* 0.1 or *G*_*b*_ *<* − 0.1, and *G*_*d*_ ≤ 0.1), purple colour indicates that both developmental plasticity and anticipatory effects evolved (in addition *G*_*e*_ *>* 0.1), green colour indicates that increased developmental variation evolved (*G*_*h*_ *>* 0.1), and black colour that there was no major change in reaction norm slope (−0.1 ≤ *G*_*b*_ ≤ 0.1). Note the logarithmic scale for *R*.

### Anticipatory effects

Anticipatory effects mediated by epigenetic changes evolved consistently when the environment did not change too fast or too slow, 1 *< R <* 100 (0 *<* log_10_ *R <* 2), and environmental predictability was 0.4 or larger (Figure 2). Very fast environmental changes do not favour the evolution of anticipatory effects, because the environment can change between when the effects are induced in the parents and when offspring develop. Very slow environmental changes also do not favour anticipatory effects as the fitness costs of anticipatory effects do not outweigh their benefits. Finally, the environment has to be rather predictable, which is understandable, since not being able to predict what the offspring environment will be can result in maladaptive anticipatory effects. Anticipatory effects evolved together with either reversible plasticity (Figure S6) when 3.2 ≤ *R* ≤ 10 (0.5 ≤ log_10_ *R* ≤ 1), or together with developmental plasticity (Figure S7) when 10 ≤ *R* ≤ 32 (1 ≤ log10 *R* ≤ 1.5).

The probability of individuals using epigenetic modifications to mediate anticipatory effects was highest when environmental predictability was more than 0.5 and when 10 ≤ *R <* 100 (1 ≤ log_10_ *R <* 2) (Figure 3). Lower values of of *P* and *R* gave slightly lower probabilities of epigenetic modification, until they epigenetic modification no longer happened.

**Figure 3:**
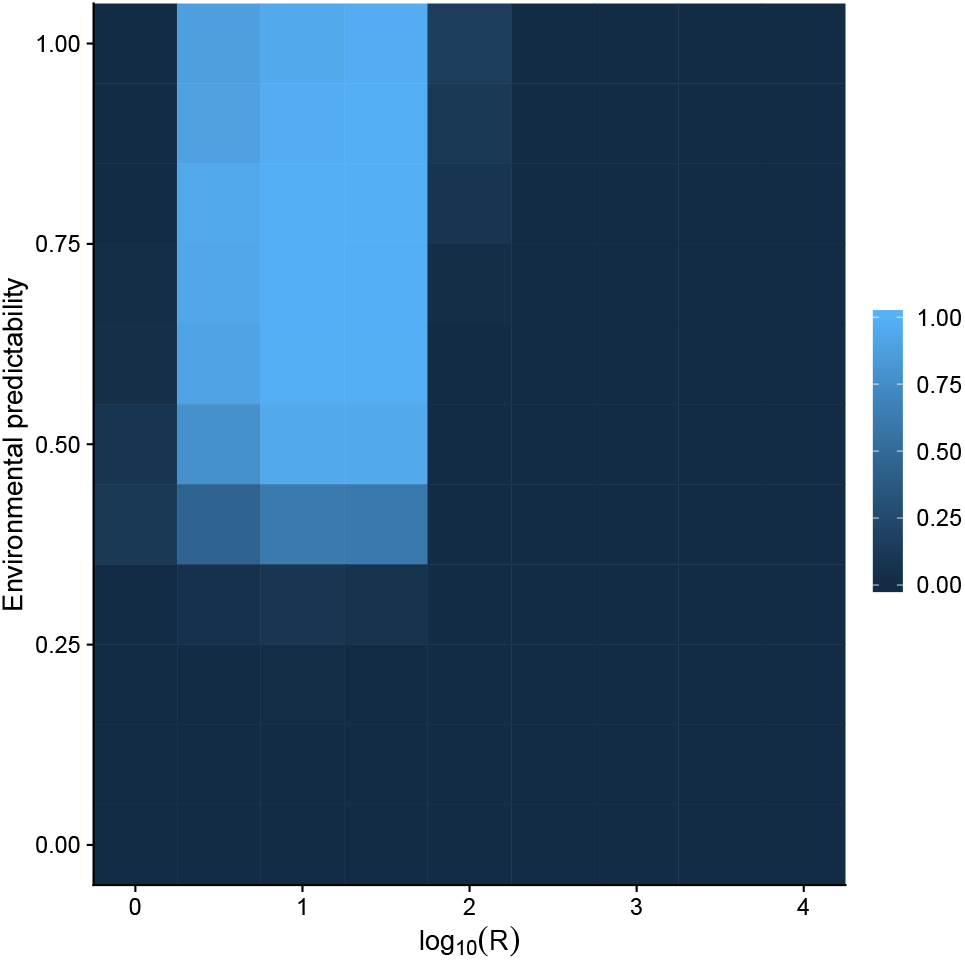
Heat map for mean probability of epigenetic modification from 100 replicate simulations with each combination of *P* and *R*. Costs of plasticity were *k*_*e*_ = *k*_*d*_ = 0.02 and *k*_*a*_ = 0.01.

Anticipatory effects increased the final mean fitness of individuals within a population. In the simulation fitness is defined as how close individual’s phenotype is to the environmental optimum, and as the environment changes population mean in fitness can change or oscillate. Therefore I used the geometric mean of the last 500 generations of mean fitness. For example, when *R* = 31.6 (log_10_ *R* = 1.5) and *P* = 0.8 the 95% quantiles for geometric mean fitness for the last 500 generations in the simulation was 0.72 [0.6, 0.74] for simulations where evolution of anticipatory effects was impossible, that is there were no loci that coded for epigenetic modification, and 0.76 [0.6, 0.77] for a simulation where anticipatory effects could freely evolve. This difference means a relative fitness of 1.05 in favour of anticipatory effects, which is more than enough to drive their evolution.

### Costs of plasticity and anticipatory effects

Fitness costs of plasticity had a large effect on what strategy the population evolved in a particular environment. When fitness costs were assumed to be zero, reversible phenotypic plasticity with anticipatory effects was very common throuhout the parameter space (Figure 4A). Clearly if there is no cost of anticipatory effects or plasticity, they should be very common, even if environmental cues predict the environment poorly. In figure 4A there is a strip at *R* = 316.2 (log_10_ *R* = 2.5) where no anticipatory effects evolve but they seem to evolve at higher values of *R*. This can actually be explained by dynamic alternation of plasticity and genetic assimilation. When *R >* 100 (log_10_ *R >* 2) and *P >* 0.3 the environment changes slowly, so that the population tracks the environment mainly by genetic adaptation. However, when there are no costs of plasticity, the population evolves transient plasticity and anticipatory effects during those periods that the environment is moving to one direction. Then when the environmental change slows, the slope of the reaction norms evolves towards zero, only to depart from zero again when the environment start moving again (Figure S8). These types of dynamics have been observed before in models of plasticity and maternal effects, where there is an extraordinary environmental change that the population has to adapt to (Lande, 2009; Hoyle & Ezard, 2012). Plastic and anticipatory effects help speed up adaptation. So this region of parameter space is characterized by transient anticipatory effects and genetic assimilation. Evolutionary changes tend to happen in reaction norm slope, the probability of epigenetic modification changes occasionally, but evolves to moderate or high value. The fitness benefits of these transient changes in plasticity are small, as any costs of plasticity prevented this type of dynamic from occurring (Figure 4). The strip at *R* = 316.2 (log_10_ *R* = 2.5) is due to environmental fluctuation being in a phase where genetic assimilation has happened and slope is zero in these populations (Figure S8A).

**Figure 4:**
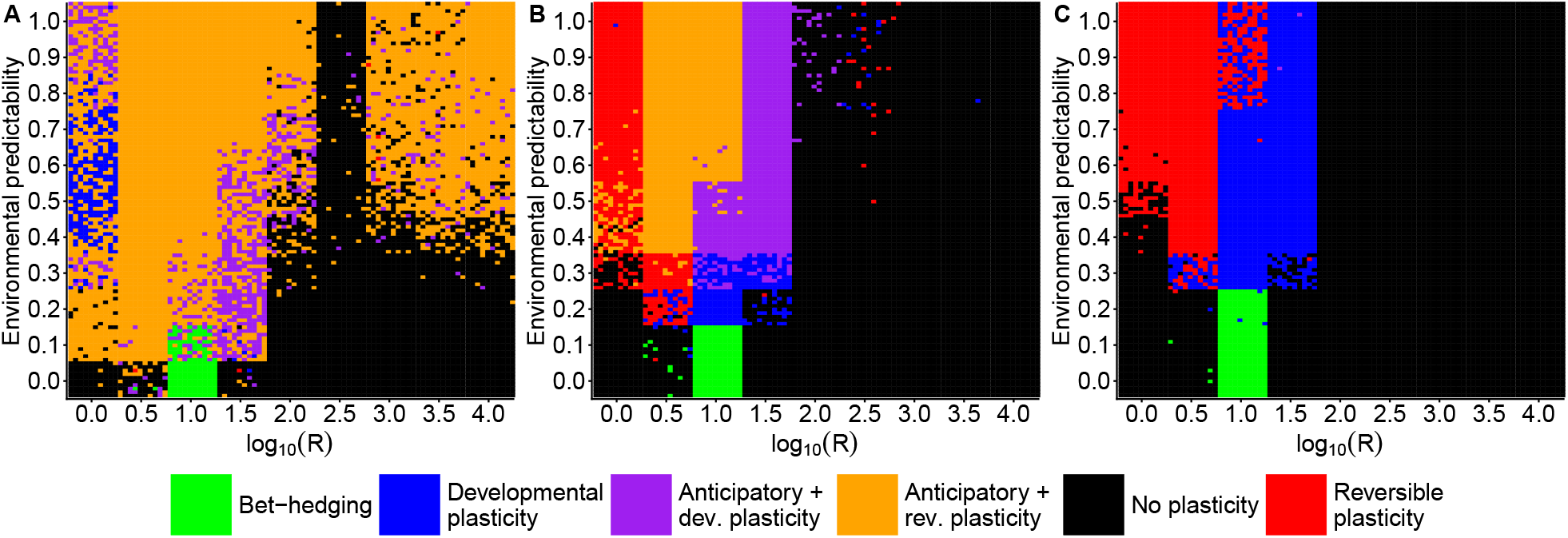
Simulation results with different costs for phenotypic plasticity and anticipatory effects. For each parameter combination of *P* and *R*, 100 replicate simulations are represented as coloured tiles, with colors corresponding to different strategies evolved in that replicate. Costs are: A) *k*_*e*_ = *k*_*d*_ = *k*_*a*_ = 0 B) *k*_*e*_ = *k*_*d*_ = 0.02 and *k*_*a*_ = 0.01 C) *k*_*e*_ = *k*_*d*_ = 0.04 and *k*_*a*_ = 0.02.

However, there was a region of parameter space where anticipatory effects did not evolve even without costs. If *P* = 0 anticipatory effects did not evolve, or if *R* ≥ 100 (log_10_ *R* ≥ 2) and *P* ≤ 0.3 no plasticity or anticipatory effects evolved (Figure 4A). In this region of parameter space anticipatory effects are likely to be either neutral or possibly deleterious as preparing offspring to a wrong environment can be maladaptive. Increasing fitness costs of plasticity and epigenetic modification reduces the range where anticipatory effects evolve (Figure 4B) and when both plasticity and epigenetic modification have high enough costs, only reversible or developmental plasticity evolves (Figure 4C).

### Influence of genetic architecture

To test whether the results were robust to genetic architecture of the phenotype, I ran a sensitivity analysis where parameters describing the genetic architecture of the population were randomly selected. The results of sensitivity analysis show that genetic architecture does influence to outcome of the simulations, but similar broad patterns are still recovered as with the initial simulations (Figure 5). Anticipatory effects do evolve, but now higher environmental predictability is generally required, *P* ≥ 0.6. Moreover, anticipatory effects now don’t evolve as certainly as in the original simulation even if plasticity evolves when 3.2 ≤ *R* ≤ 31.6 (0.5 ≤ log_10_ *R* ≤ 1.5) (Figure 5). Since the fitness benefits of anticipatory effects are smaller than within generation plasticity, sometimes genetic architecture does not allow fine-tuning of the anticipatory effects, which is seen especially in lower values of environmental predictability where lower probability of epigenetic modification evolves (Figure 3). In some cases the population goes extinct because there is not enough genetic variation for adaptation, but this was not common as only 51 extinction events were observed which is 0.5% of the total simulations. In the sensitivity analyses the parameter space where increased developmental variation or a bet-hedging type stategy could evolve was actually larger than in the original simulations. If the environmental component of random variation was small or when other strategies were more restricted there was more opportunity to sometimes evolve increased developmental variation (Figure 5).

**Figure 5:**
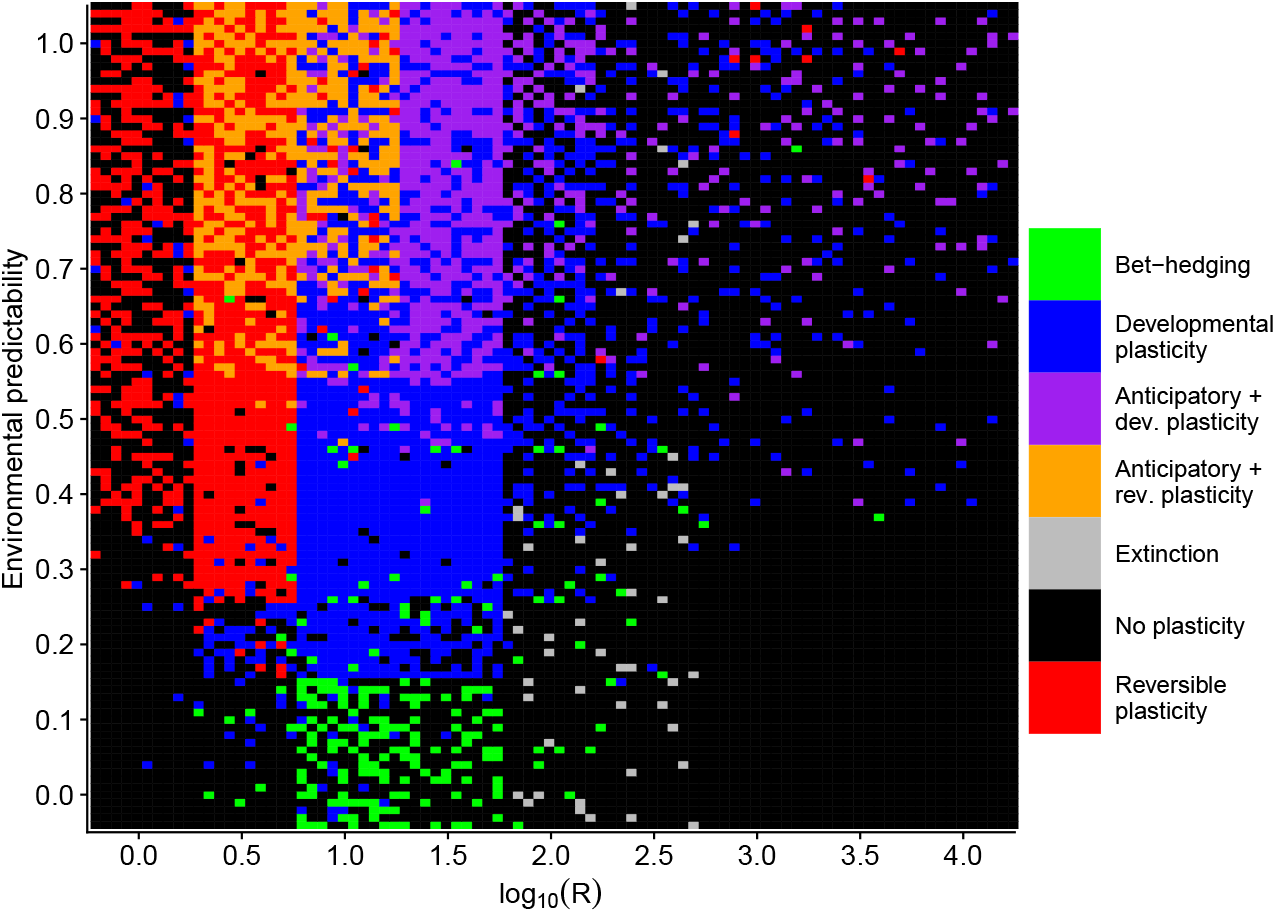
Sensitivity of results to genetic architecture. 100 replicate simulations were run for each combination of *R* and *P*, where parameter values for the random environmental effects, number of loci, number of chromosomes, mutation rate, and mutational effects were drawn randomly. Costs of plasticity were *k*_*e*_ = *k*_*d*_ = 0.02 and *k*_*a*_ = 0.01. Coloured tiles show which strategy evolved in the population or if it went extinct.

To investigate further the influence of specific parameters of genetic architecture on the evolution of anticipatory effects I ran simulations changing the number of loci that were responsible for the phenotypic traits or the standard deviation of mutational effects. The number of loci had the strongest effect on the evolution of reversible plasticity. When the slope of the reaction norm and probability of adjustment were both determined by only one locus, evolution of reversible plasticity was prevented in an environment that completed its cycle in every generation, *R* = 1 (log_10_ *R* = Anticipatory effects evolved when *P* ≥ 0.6 and *R* ≥ 3.2 (log_10_ *R* ≥ 0.5) (Figure 6). Moreover, increased developmental variability evolved when environmental predictability was low, *P* ≤ 0.1, and rate of environmental change was intermediate, 10 ≤ *R* ≥ 32 (1 ≤ log_10_ *R* ≥ 1.5). When the number of loci was increased to 5 reversible plasticity evolved also in environments that fluctuated every generation (Figure 6B). Further increasing the number of loci to 20 allowed reversible plasticity to evolve in more unpredictable environments (Figure 6C). Increasing the number of loci did allow anticipatory effects to evolve when *P* = 0.5 in cases where developmental plasticity had evolved. In general the number of loci had only small effect on the evolution of anticipatory effects.

**Figure 6:**
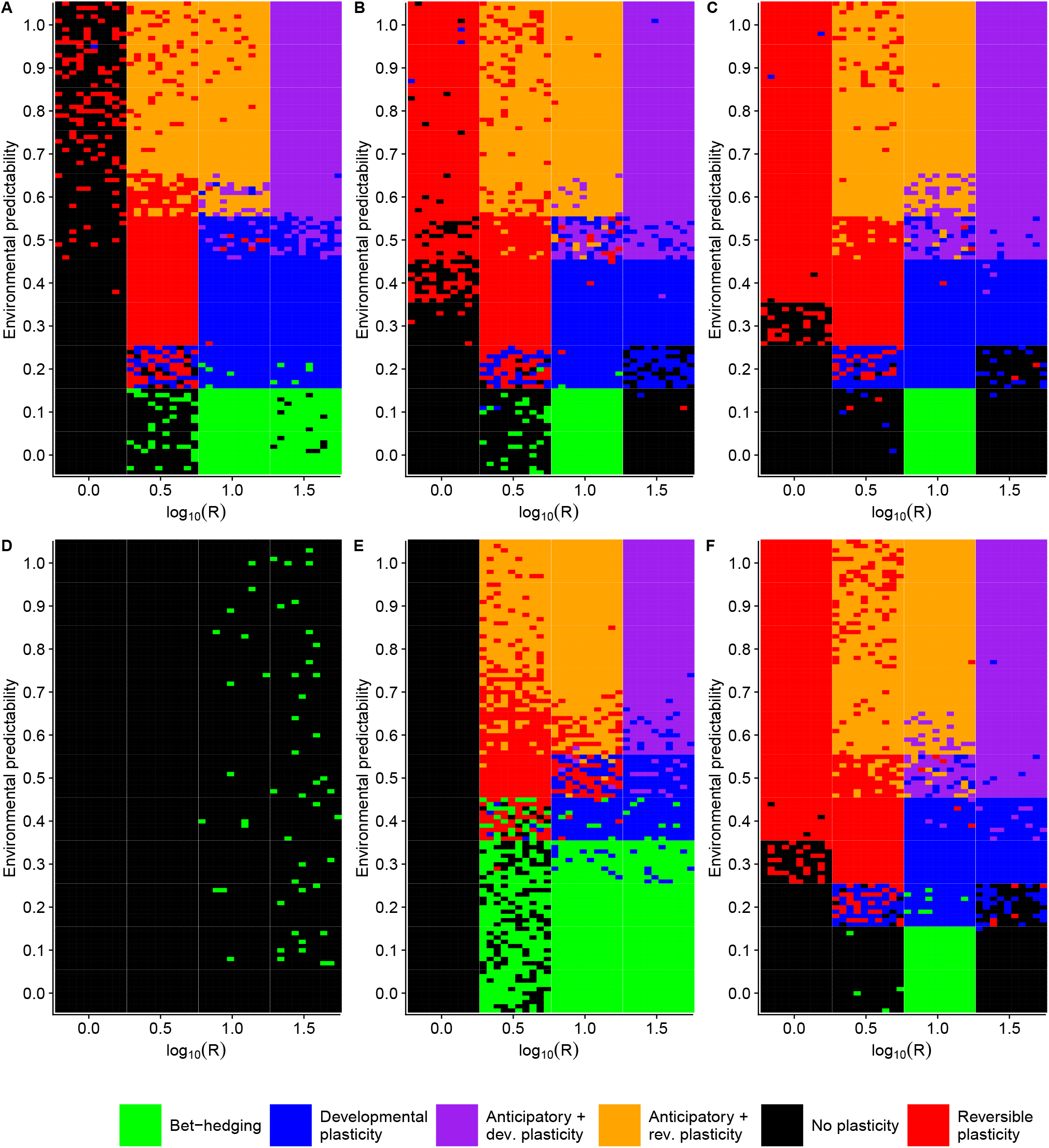
Effects of number of loci (A to C) and standard deviation of mutational effects (D to F). 100 replicate simulations were run for each combination of *R* and *P*, but only those values of *R* were used where I had previously observed the evolution of plasticity when plasticity had a cost. Cost of plasticity were *k*_*e*_ = *k*_*d*_ = 0.02 and *k*_*a*_ = 0.01. Coloured tiles show which strategy evolved in the population. A) *l*_*a*_ = *l*_*b*_ = *l*_*d*_ = *l*_*e*_ = *l*_*h*_ = 1 B) *l*_*a*_ = *l*_*b*_ = *l*_*d*_ = *l*_*e*_ = *l*_*h*_ = 5 C) *l*_*a*_ = *l*_*b*_ = *l*_*d*_ = *l*_*e*_ = *l*_*h*_ = 20 D) *σ*_*α*_ = 0.01 E) *σ*_*α*_ = 0.1 F) *σ*_*α*_ = 2.

Next I investigated the role of mutational effects in the evolution of anticipatory effects. I ran simulations with different standard deviation for mutational effects. When *σ*_*α*_ = 0.01 no plasticity of any kind evolved (Figure 6D). When this was increased to *σ*_*α*_ = 0.1, anticipatory effects did evolve when *P* ≥ 0.6 and *R* ≥ 3.2 (log_10_ *R* ≥ 0.5) (Figure 6E). However, no reversible plasticity evolved in simulation when *R* = 1 (log_10_ *R* = 0). The reason plasticity did not evolve in these conditions was that rate of evolution was too slow relative to the rate of environmental change due to the small mutational effects. When *σ*_*α*_ = 0.01 and *R* is small there is no evolution of plasticity (Figure S9A), but when *R* is large, *R* = 3162.3 (log_10_ *R* = 3.5), so that the population evolves a strategy to track the environmental optimum by genetic adaptation, there is enough time to reach a stable state for the genotypic values of the intercept (Figure S9B). Therefore, it is not the case that there no time to reach equilibrium during the 1500 generations, but that selection pressure changes too fast relative to the slow rate of evolution that would be enough time for the small effects to build up. This is also seen for *σ*_*α*_ = 0.1, but in less extreme form, as only *R* = 1 (log_10_ *R* = 0) is too fast to prevent adaptation (Figure S9C), while *R* = 3.2 (log_10_ *R* = 0.5) already allows stable states to be reached (Figure S9D). Furthermore, increased developmental variation evolved in up to *P* ≤ 0.3 and 3.2 ≤ *R* ≥ 32 (0.5 ≤ log_10_ *R* ≥ 1.5) (Figure 6E), which represents much larger parameter space than with larger mutational effects (Figure 6F). When mutational effects were double the default value, *σ*_*α*_ = 2, resersible plasticity evolved as normal, and anticipatory effects evolved also when predictability of the environment was slightly lower *P* = 0.5 in cases where developmental plasticity had evolved. In the case where anticipatory effects did not evolve, substituting the parameter values used in the simulation gives a mutational variance of 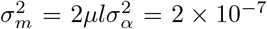 and mutational heritability of 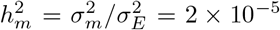. These values are orders of magnitude lower than values observed empirically for many phenotypic traits. In summary, anticipatory effects evolved unless mutational effects were very small and it seems that genetic architecture is unlikely to completely prevent evolution of anticipatory effects if they are ecologically favoured.

## Discussion

The results reported here largely agree with previous modeling results concerning maternal effects in phenotypic memory type of models, in that anticipatory effects can evolve as long as environmental cues can predict the environment of the offspring to some degree (Hoyle & Ezard, 2012; Ezard et al., 2014; Kuijper & Hoyle, 2015; Kuijper & Johnstone, 2015; Leimar & McNamara, 2015). The predictive power of environmental cues does not have to be perfect as anticipatory effects still evolved when environmental predictability was rather low. The costs of anticipatory effects influence the threshold of minimum predictability, since when there were no costs for any type of plasticity, anticipatory effects evolved even with minimal predictability. As usefulness of environmental cues in predicting the offspring environment decreases, less and less fitness benefit can be gained from anticipatory effects, and therefore only such anticipatory effects that have very low costs or no costs at all can evolve in rather unpredictable environments. Only a predictability of zero meant that there was not evolution of any kind of anticipatory effects.

Anticipatory effects evolved only when frequency of environmental fluctuations was slow enough that cues perceived by the parents still meant something when the offspring were developing, but fast enough there could be differences between the parent and offspring environments. So, if environmental change was slow enough, there is little point of paying the potential costs of anticipatory effects if the fitness benefit gained from such an effect is small. In contrast to intermediate fluctuation frequencies favouring anticipatory effects, in certain models phenotypic memory is favoured in very fast fluctuations (Jablonka et al., 1995; Lachmann & Jablonka, 1996). However, this can be explained regarding how these models were defined, in Jablonka et al. (1995) and Lachmann & Jablonka (1996) the environment fluctuated between two discrete states. Thus it makes sense that faster fluctuations might favour anticipatory effects in these models. Other models of environmental epigenetic induction have substituted temporal environmental fluctuations for spatial heterogeneity in the environment, and if one considers temporal and spatial variation as analogous, at least qualitatively similar results emerge that migration between populations has to be moderate for anticipatory effects to evolve (Greenspoon & Spencer, 2018).

The evolution of anticipatory effects is obviously dependent the magnitude of the fitness benefit they give relative to their costs. In the simulation when anticipatory effects had high costs, *k*_*e*_ = 0.04, the evolution of anticipatory effects was completely prevented. Understandably so, as in this model anticipatory effects gave only a moderate fitness benefit overall and within generation plasticity was much more beneficial. This phenomenon has also been observed by Kuijper & Hoyle (2015), as using direct information about the current environment is always going to be better than using incomplete information about future environment. Therefore, in real populations one could expect anticipatory effects to be smaller in magnitude or rarer than within generation plastic effects, this smaller fitness benefit possibly explains why anticipatory effects seem to be generally weak (Uller et al., 2013). Most theory about the evolution of plasticity in general assumes that there are some costs of plasticity (Callahan et al., 2008). However, while some studies have detected costs of plasticity (Relyea, 2002; Dechaine et al., 2007), in general costs of plasticity have been notoriously difficult to document in nature and we have a poor idea if we expect costs of anticipatory effects to be generally the same or somehow different. As fitness benefits of anticipatory effects are smaller than for within generation plasticity, it may be that costs of anticipatory effects are an important component limiting their evolution.

Certain organisms seem to exhibit a lot of anticipatory effects, such as nematodes (Baugh & Day, 2020). This may reflect an experimental bias, as many people are studying these model organisms. However, another explanation is that there is something in their ecology that favors the evolution of anticipatory effects. Environments change at different rates from the perspective of different organisms since generation times can be vastly different. What are fast environment changes for long-lived multicellular organisms can be rather slow changes from the perspective of unicellular microbes. It may be, perhaps somewhat counterintuitively, that anticipatory effects can be more common in smaller organisms that are short-lived. Moreover, the implication also is that organisms which have a long period between zygote formation and emergence of live offspring, such as mammals with long gestation periods, are expected to have less anticipatory effects than organisms where there is a shorter time between zygote formation and emergence. The longer the time between setting the epigenetic marks in gametes and offspring birth, the smaller the correlation between the environment of the offspring and the environment of the parent. In plants it is known that anticipatory effects are sometimes transmitted through seed but not by pollen (Herman et al., 2014; Wibowo et al., 2016), presumably because pollen can disperse over much longer distances, diminishing the correlation between parent and offspring environment. Unequal resetting of epigenetic marks in male and female gametes should apply as well to species where the time of gamete production and the time of fertilization are decoupled, such as certain insects, reptiles, and birds that are capable of storing sperm for long periods of time (Boomsma et al., 2005; Holt & Fazeli, 2016).

Another important ecological question is how predictable real environments are from the perspective of organisms. Are there environmental cues that organisms can potentially use to predict future environments? The most obvious cues are related to changes in day length which does reliably predict seasons in high and low latitudes. Many environmental variables can also function as cues themselves, such as temperature. For example, high temperatures today can predict that temperature will also be high in the short term. Furthermore, there are some indications that many environmental variables are autocorrelated in nature (Halley, 1996), and that this autocorrelation often tends to be positive (Vasseur & Yodzis, 2004; Ruokolainen et al., 2009), meaning that temporally close observations tend to be similar. Moreover, there seem to be differences in terrestrial and marine ecosystems; autocorrelation in terrestrial systems seems to be closer to zero, while marine ecosystems tend to have more positive autocorrelations (Vasseur & Yodzis, 2004; Ruokolainen et al., 2009). It will be interesting to see if anticipatory effects are more prevalent in marine organisms. Nevertheless, environments in nature tend to be at least somewhat autocorrelated, so there should be some opportunity for evolution of anticipatory effects.

In some empirical studies of epigenetically mediated anticipatory effects, the effects induced by the environment last for several generations. This has been particularly observed in nematodes (Baugh & Day, 2020). The question remains whether these multigenerational effects are adaptive in nature, or rather a consequence of some developmental constraint? In the model used in this study it is difficult to see how multigenerational effects could be adaptive, as this would require the environment of the grandparents to predict the environment of the offspring better than the environment of their parents. In the model epigenetic modifications were reset at every generation, but relaxing this assumption would likely not change the results, as it makes sense for the parents to overwrite any previous epigenetic marks using the current environmental cue to maximize the correlation between their environment and the offspring environment. This assumption of course depends on the mechanistic details on how the epigenetic modification system works, and if there is asymmetry in the way the epigenetic mark can influence the phenotype this could potentially change the results. For example, if the epigenetic change induces some antipredator defences and predation risk remains elevated for several generations even in the absence of the initial environmental cue about predator presence, then multigenerational effects that maintain antipredator defences for several generation could be adaptive. A model investigating the incomplete resetting of epigenetic marks found that incomplete resetting was favoured when the environment changed infrequently and environmental cues were unreliable (Uller et al., 2015). In the model studied here, costs of anticipatory effects generally prevented their evolution in these kinds of conditions. Therefore, it seems that multigenerational effects will be dependent on very specific ecological conditions and are more likely to occur in organisms with very short generation times.

## Conclusions

Simulations with a genetically explicit model generally support many of the results obtained previously: anticipatory effects can evolve when the environment changes at moderate speeds relative to generation time, and there is some environmental cue that can be used to predict the environment. However, if anticipatory effects have fitness costs, these costs limit their evolution only to situations where environmental cues are reliable. If fitness costs are very low, anticipatory effects also help genetic adaptation to large environmental changes, in a manner similar to within generation plasticity and maternal effects. In terms of novel results, this model shows that the evolution of anticipatory effects is quite robust to different genetic architectures, although genetic architecture can cause the need for environmental cues to be very reliable. Unless mutational effects were very small, anticipatory effects could readily evolve. In order to test assumptions of the model, future empirical studies could benefit from characterizing the predictability of environmental cues or the autocorrelation structure of the environmental variable that induces the possible anticipatory effects. The fitness costs of anticipatory effects remain another puzzle. It has been difficult to determine fitness costs of within generation plasticity, so getting estimates of costs of anticipatory effects will undoubtedly be challenging. The model presented here could be tested experimentally by evolving eukaryotic microbes in environments that fluctuate at different frequencies and investigating whether the populations evolve plastic or anticipatory responses mediated by epigenetic changes.

## Acknowledgments

This study was funded by an Academy of Finland Research Fellowship (no. 321584). I thank the Finnish CSC-IT Center for Science Ltd. for providing computational resources.

## Supplementary Information

### Supplementary Figures

**Figure S1:**
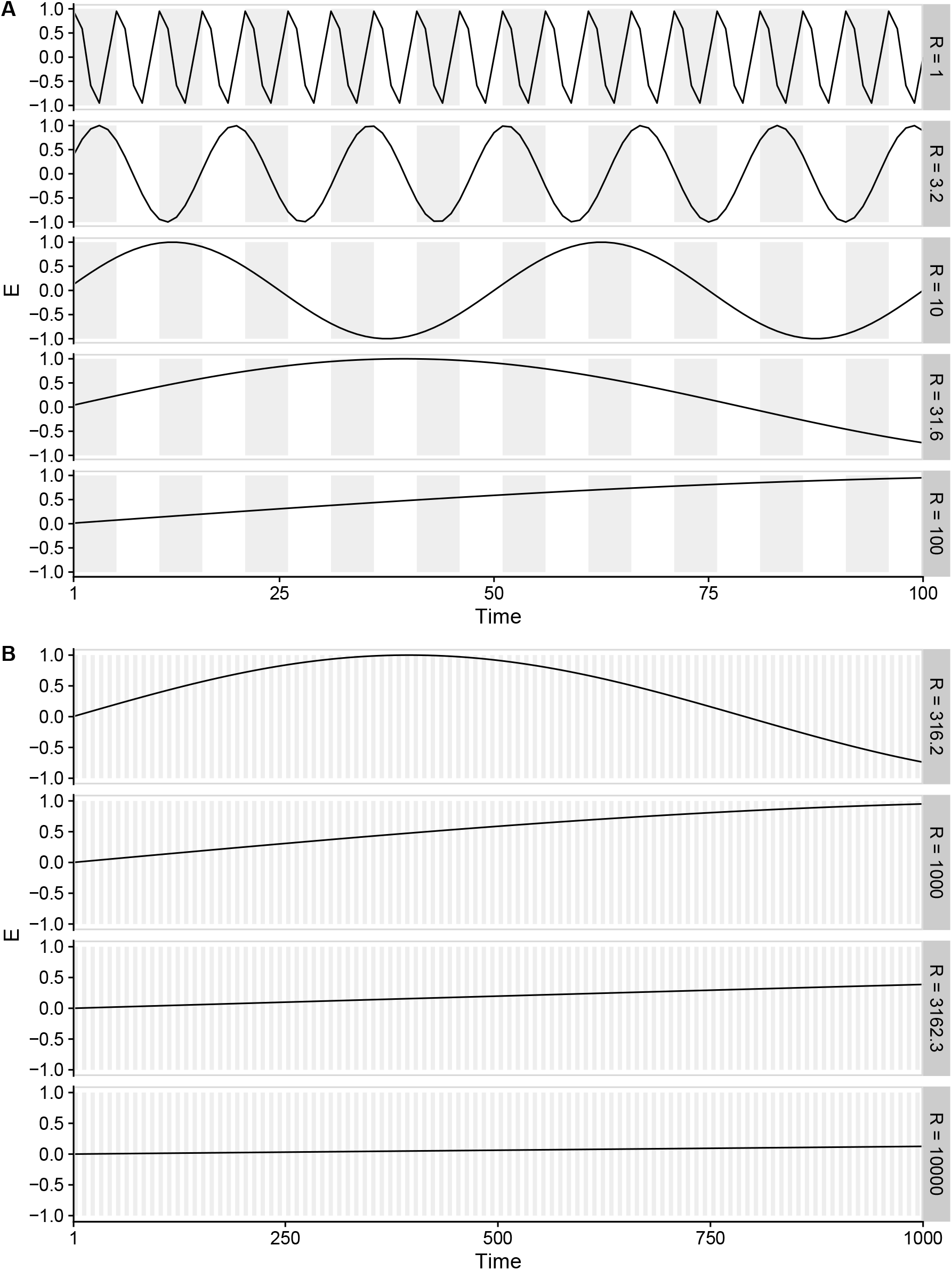
Illustration of how the environment fluctuates with different values of *R* used in the simulations. To illustrate different rates of environmental change, slower fluctuations are plotted on a different scale. A) Twenty generations are plotted. B) Two hundred generations are plotted. Each generation is five timesteps long, as shown by alternation of shading.

**Figure S2:**
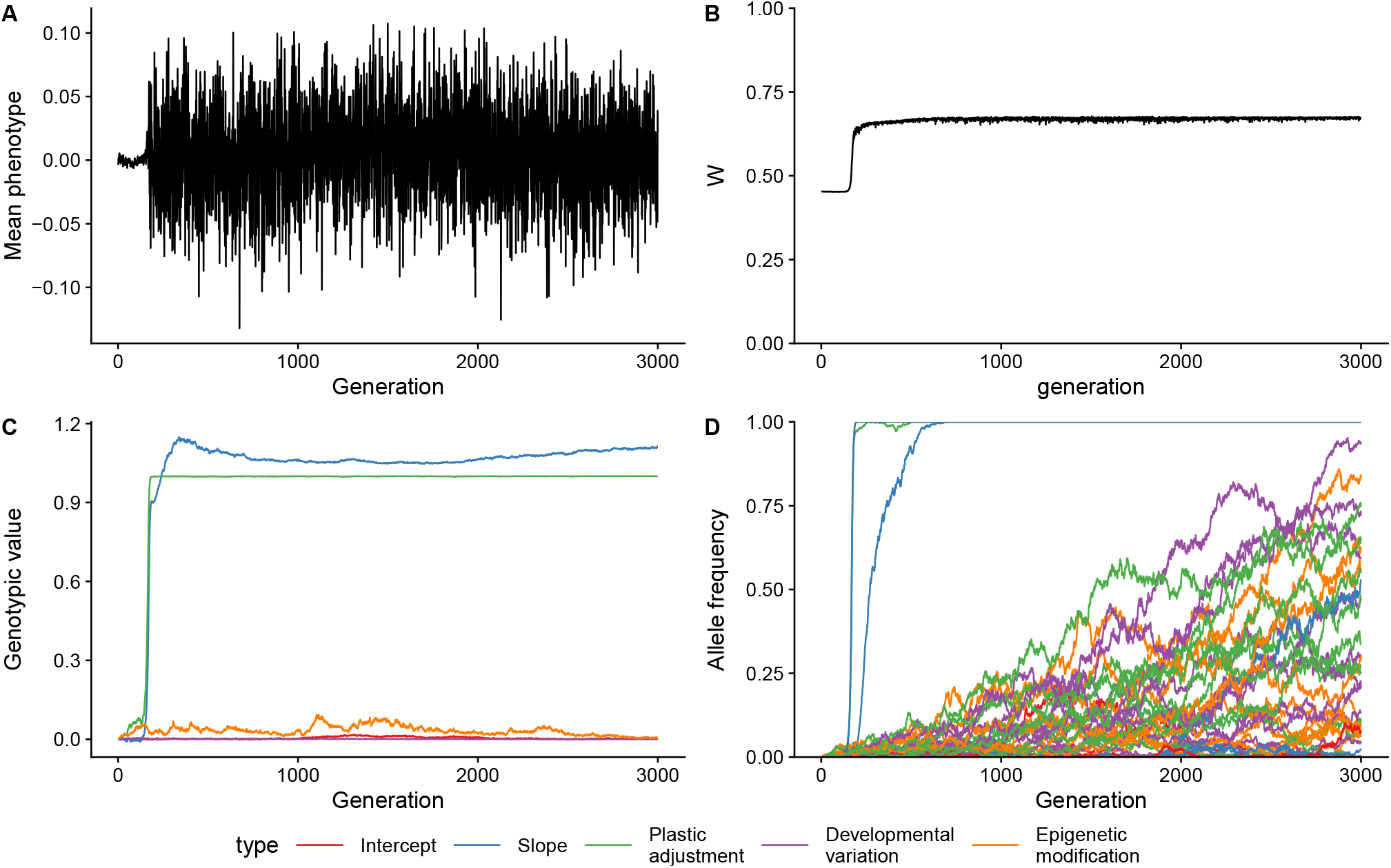
An example of how a single population evolves a strategy of reversible phenotypic plasticity. Simulation parameters were: *R* = 1, *P* = 0.9, *k*_*d*_ = *k*_*e*_ = 0.02, and *k*_*a*_ = 0.01. A) Phenotypic mean of the population for each generation. B) Population mean fitness for each generation. C) Genotypic values for reaction norm intercept, slope, probability of plastic adjustment, developmental variation, and probability of epigenetic modification for each generation. D) Allele frequencies of derived alleles at QTL controlling the genotypic values. Legend shows colours for types of genotypic values or loci.

**Figure S3:**
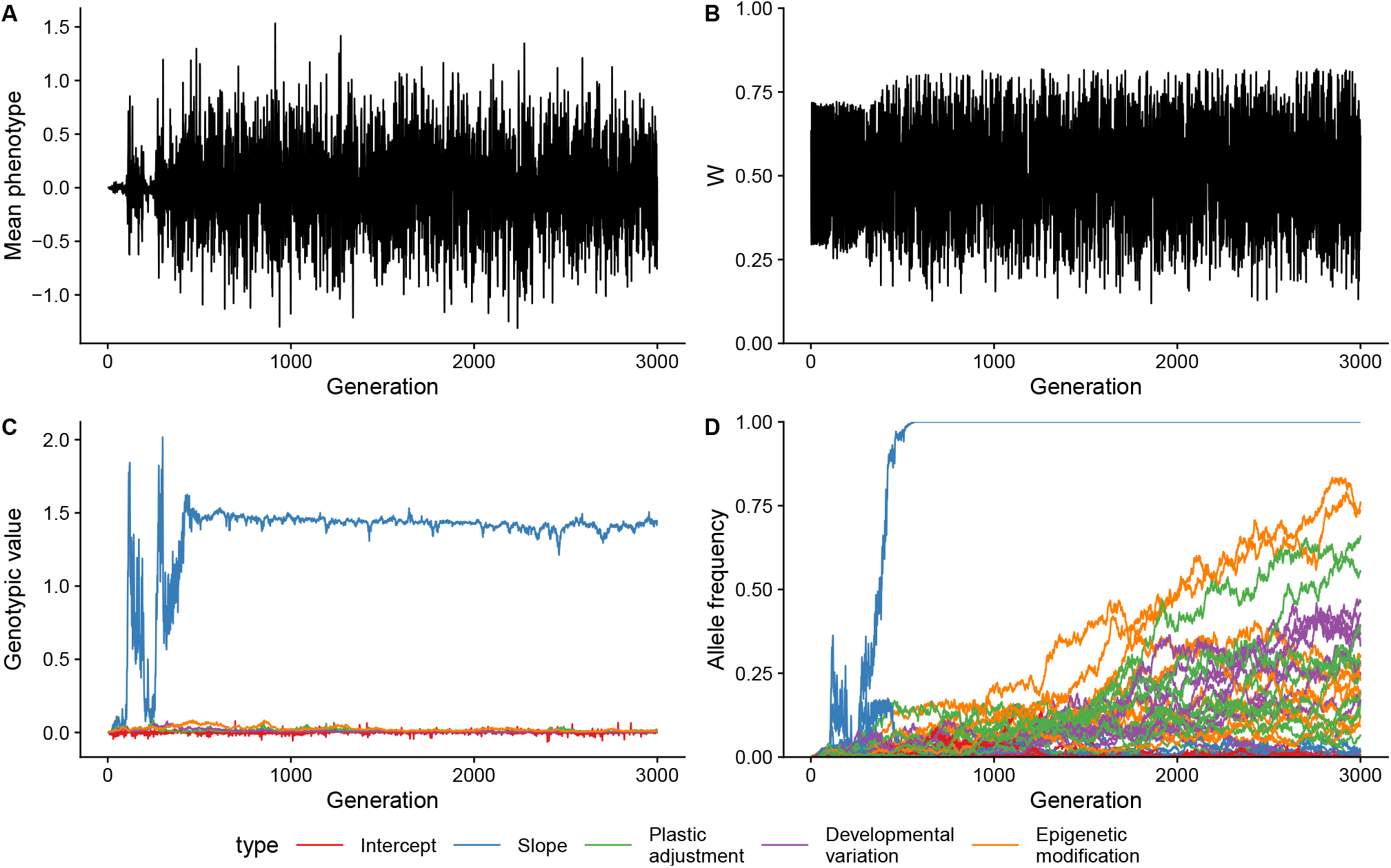
An example of how a single population evolves a strategy of developmental phenotypic plasticity. Simulation parameters were: *R* = 10, *P* = 0.2, *k*_*d*_ = *k*_*e*_ = 0.02, and *k*_*a*_ = 0.01. A) Phenotypic mean of the population for each generation. B) Population mean fitness for each generation. C) Genotypic values for reaction norm intercept, slope, probability of plastic adjustment, developmental variation, and probability of epigenetic modification for each generation. D) Allele frequencies of derived alleles at QTL controlling the genotypic values.

**Figure S4:**
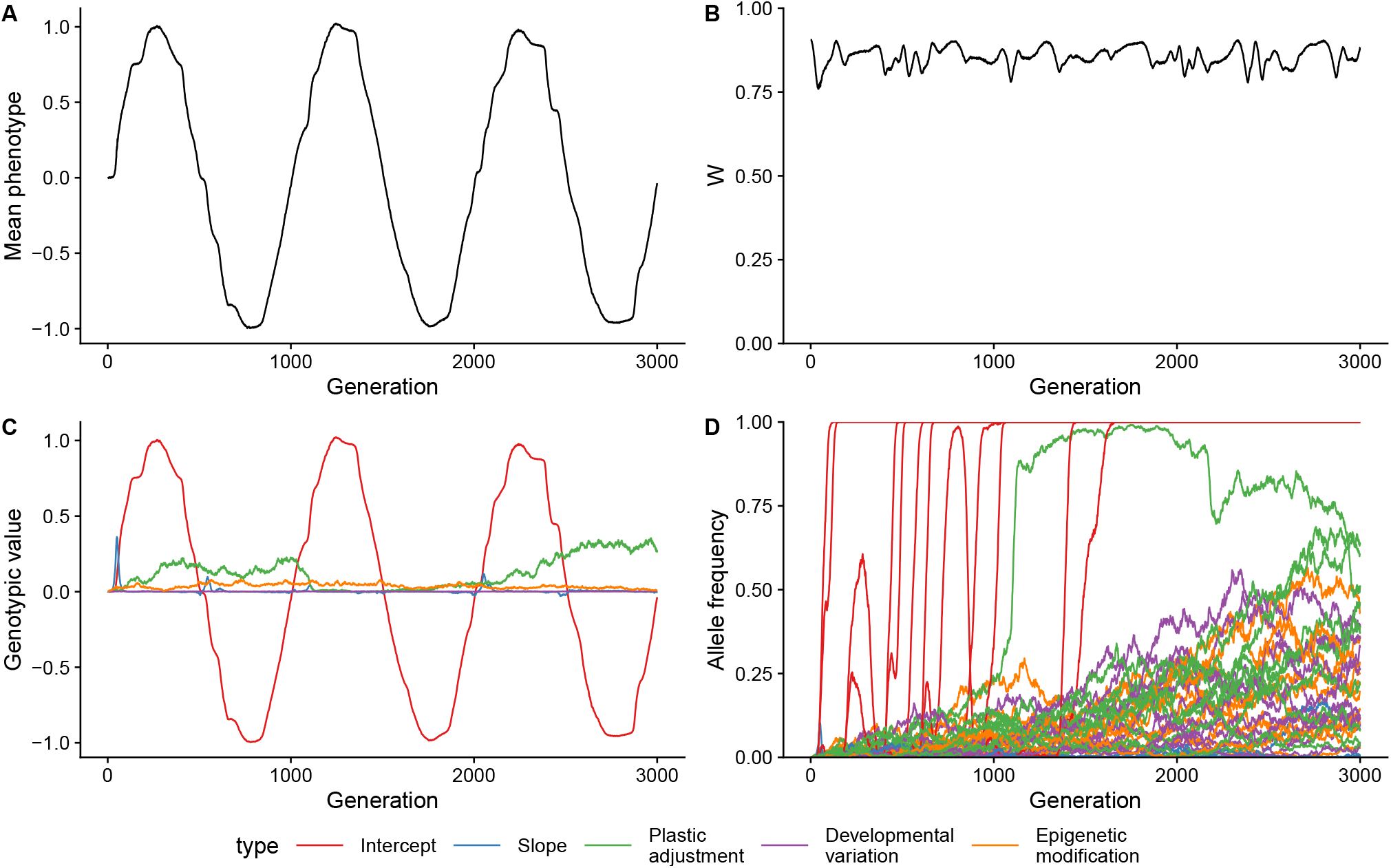
An example of how a single population evolves by tracking the environmental optimum by changing the reaction norm intercept. Simulation parameters were: *R* = 1000, *P* = 0.9, *k*_*d*_ = *k*_*e*_ = 0.02, and *k*_*a*_ = 0.01. A) Phenotypic mean of the population for each generation. B) Population mean fitness for each generation. C) Genotypic values for reaction norm intercept, slope, probability of plastic adjustment, developmental variation, and probability of epigenetic modification for each generation. D) Allele frequencies of derived alleles at QTL controlling the genotypic values.

**Figure S5:**
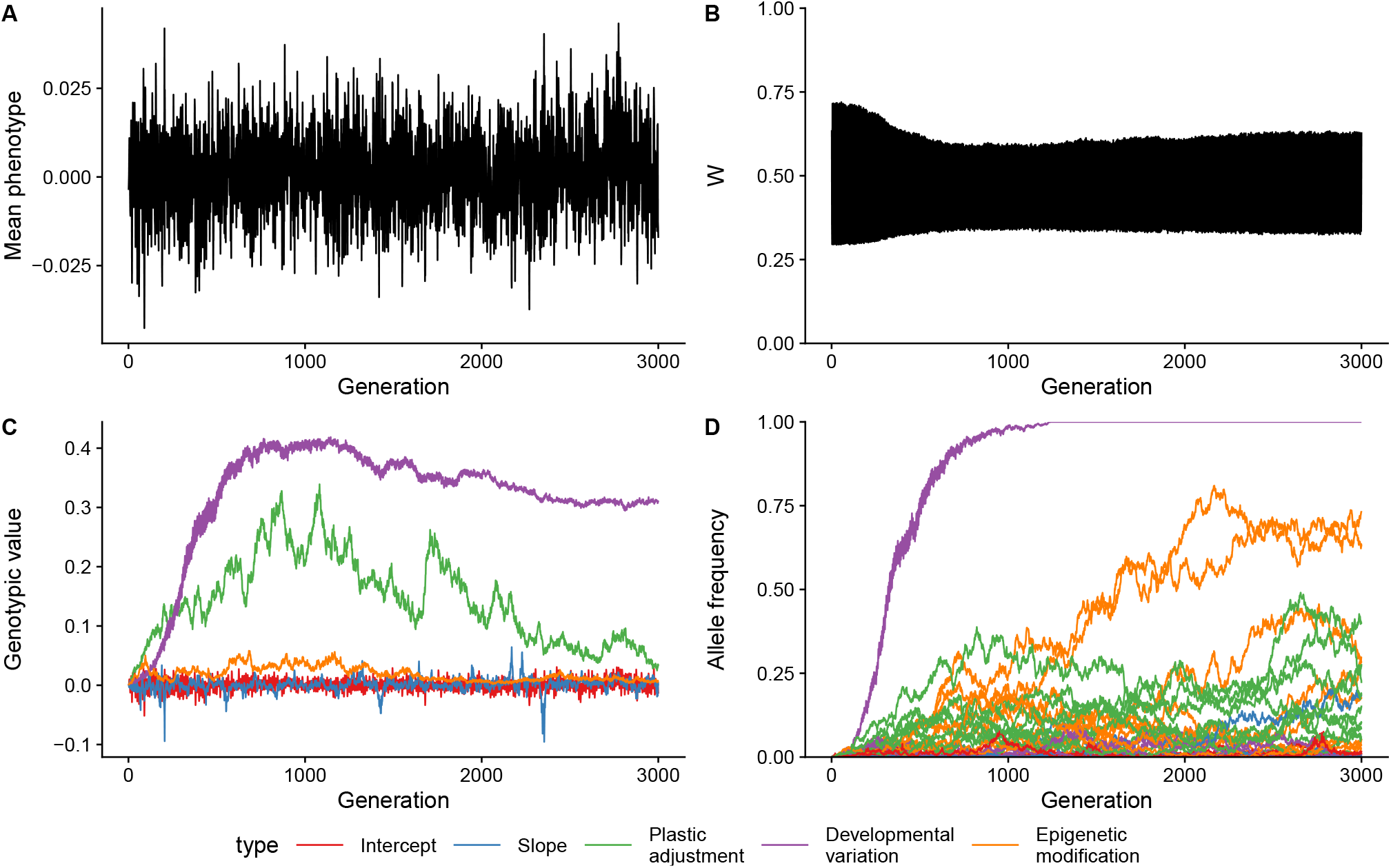
An example of how a single population evolves a strategy of diversifying bet-hedging. Simulation parameters were: *R* = 10, *P* = 0, *k*_*d*_ = *k*_*e*_ = 0.02, and *k*_*a*_ = 0.01. A) Phenotypic mean of the population for each generation. Population mean fitness for each generation. C) Genotypic values for reaction norm intercept, slope, probability of plastic adjustment, developmental variation, and probability of epigenetic modification for each generation. D) Allele frequencies of derived alleles at QTL controlling the genotypic values.

**Figure S6:**
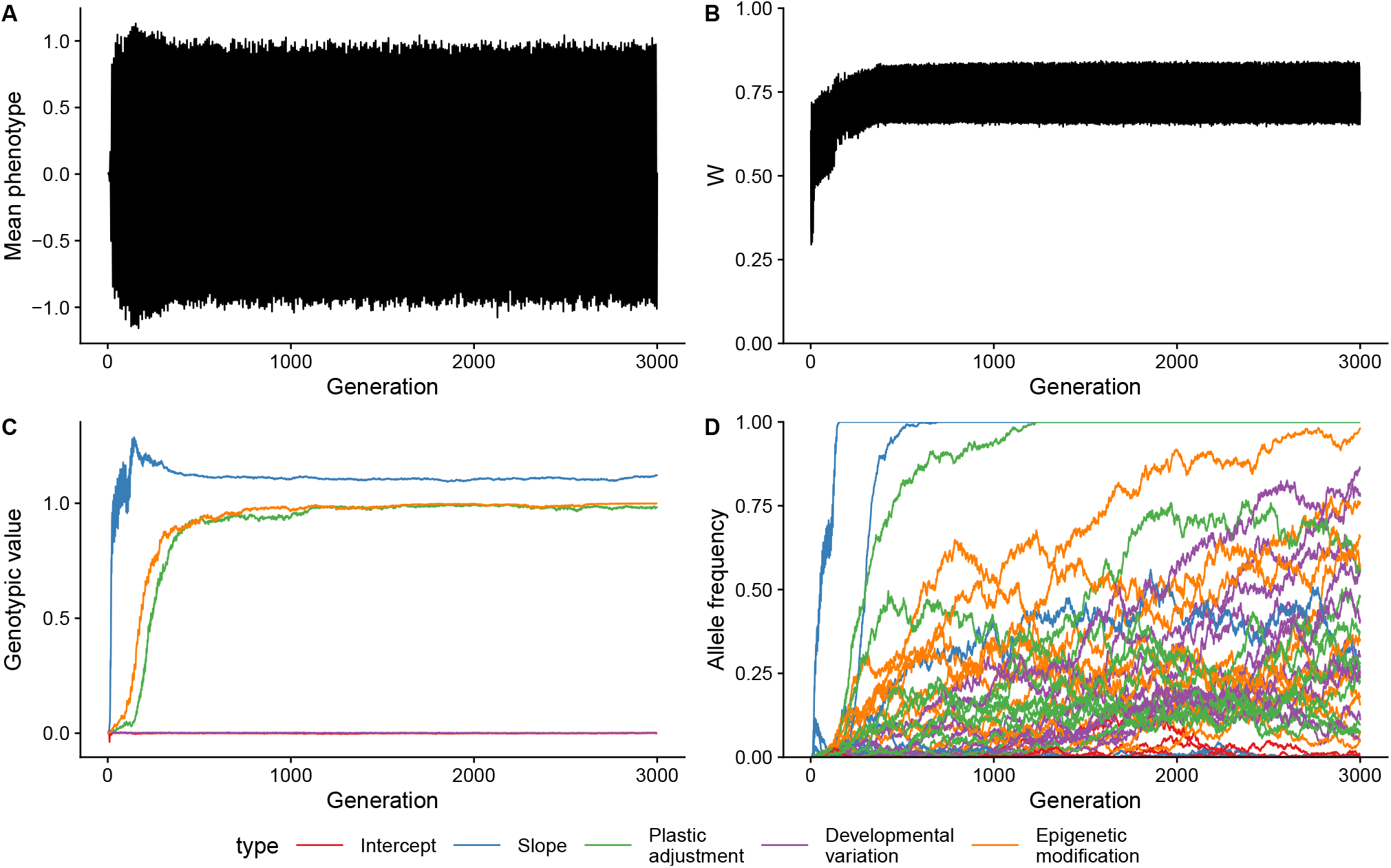
An example of how a single population evolves a strategy of reversible phenotypic plasticity and anticipatory effects via epigenetic modifications. Simulation parameters were: *R* = 10, *P* = 0.9, *k*_*d*_ = *k*_*e*_ = 0.02, and *k*_*a*_ = 0.01. A) Phenotypic mean of the population for each generation. B) Population mean fitness for each generation. C) Genotypic values for reaction norm intercept, slope, probability of plastic adjustment, developmental variation, and probability of epigenetic modification for each generation. D) Allele frequencies of derived alleles at QTL controlling the genotypic values.

**Figure S7:**
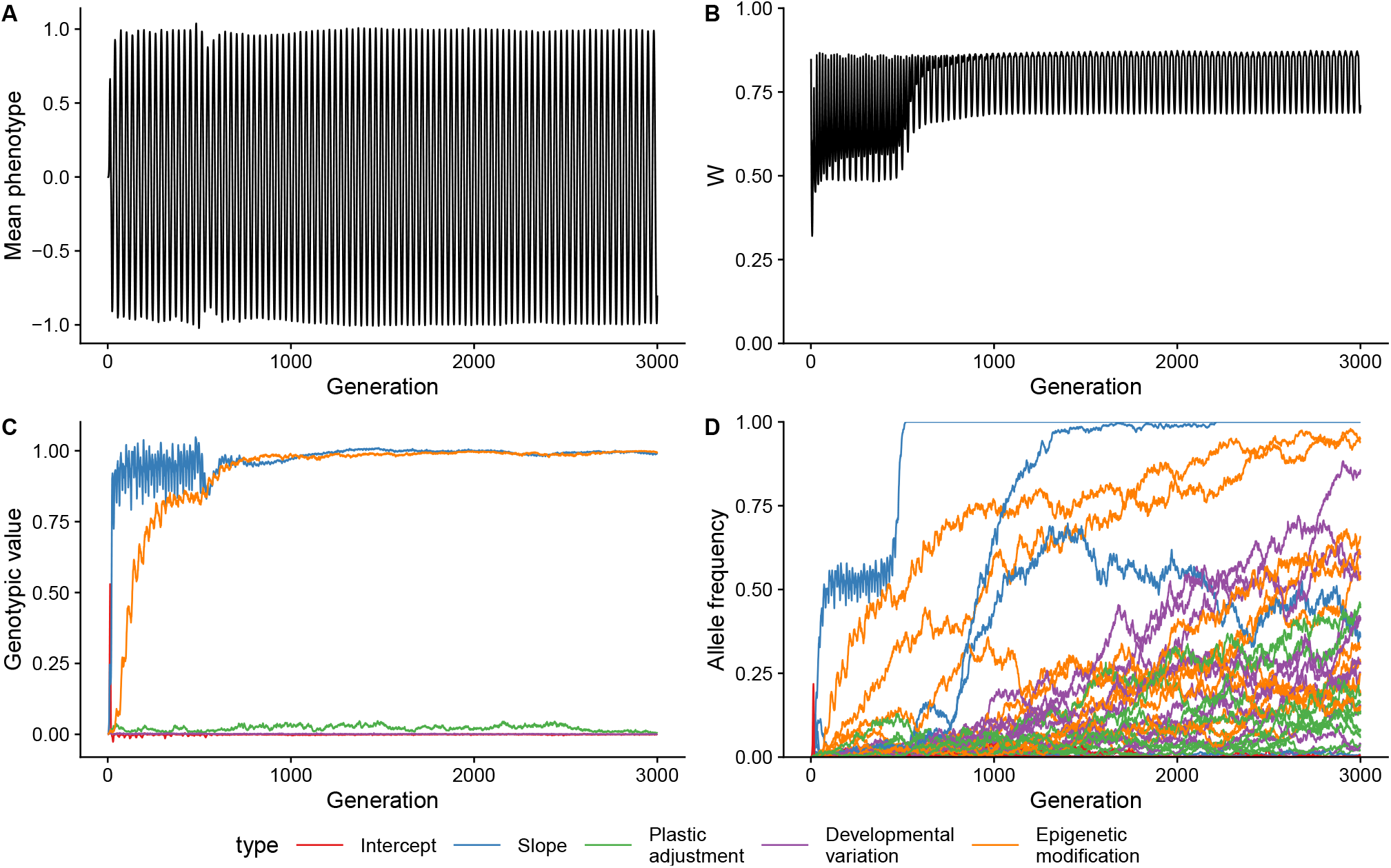
An example of how a single population evolves a strategy of developmental phenotypic plasticity and anticipatory effects via epigenetic modifications. Simulation parameters were: *R* = 31.6, *P* = 1, *k*_*d*_ = *k*_*e*_ = 0.02, and *k*_*a*_ = 0.01. A) Phenotypic mean of the population for each generation. B) Population mean fitness for each generation. C) Genotypic values for reaction norm intercept, slope, probability of plastic adjustment, developmental variation, and probability of epigenetic modification for each generation. D) Allele frequencies of derived alleles at QTL controlling the genotypic values.

**Figure S8:**
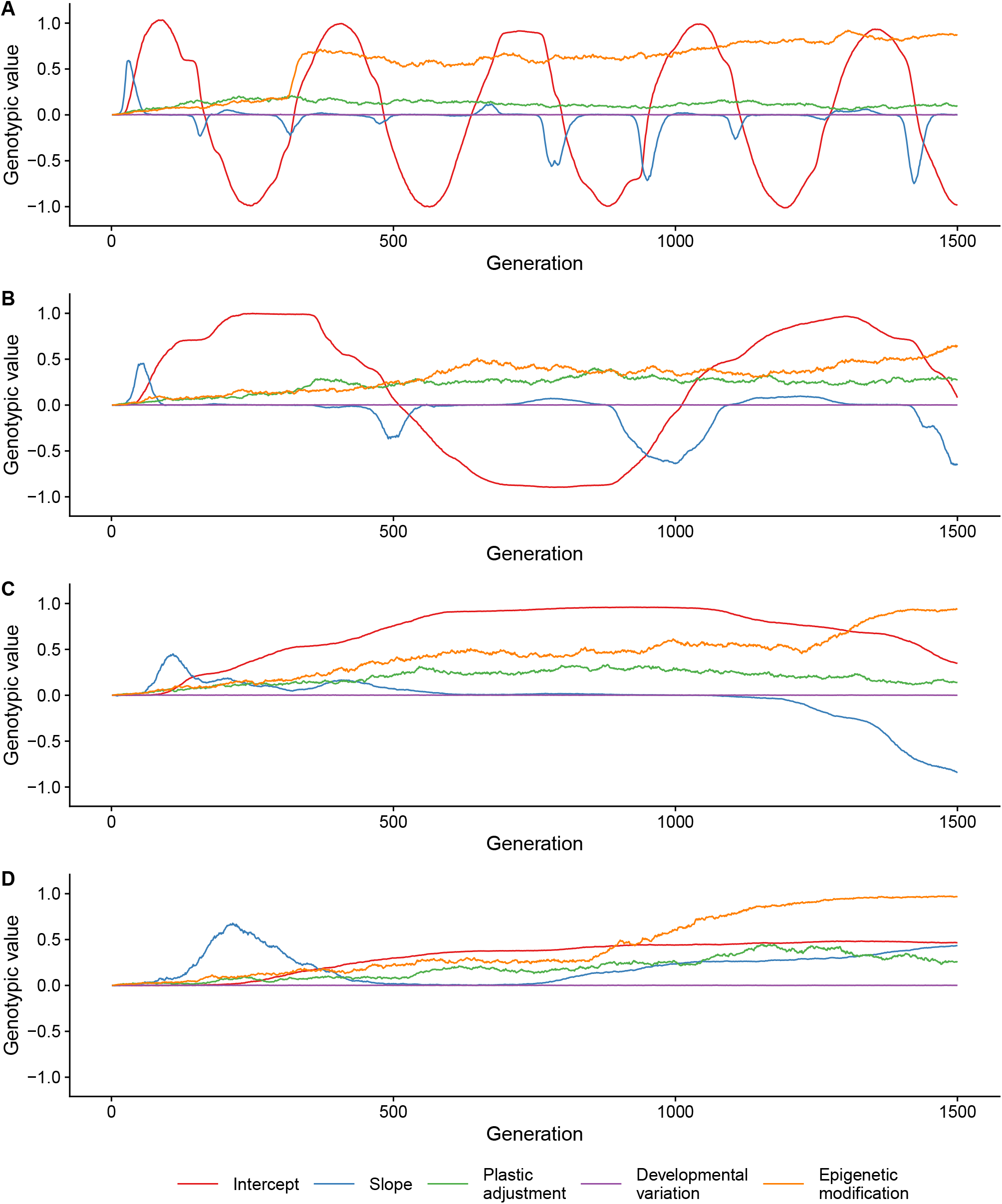
Examples of transient dynamics of plasticity and anticipatory effects punctuated by genetic assimilation. Genotypic values are shown for individual simulations runs. In all panels costs of plasticity are zero, *k*_*d*_ = *k*_*e*_ = *k*_*a*_ = 0, and *P* = 1. A) *R* = 316.2 B) *R* = 1000 C) *R* = 3162.3 D) *R* = 10000

**Figure S9:**
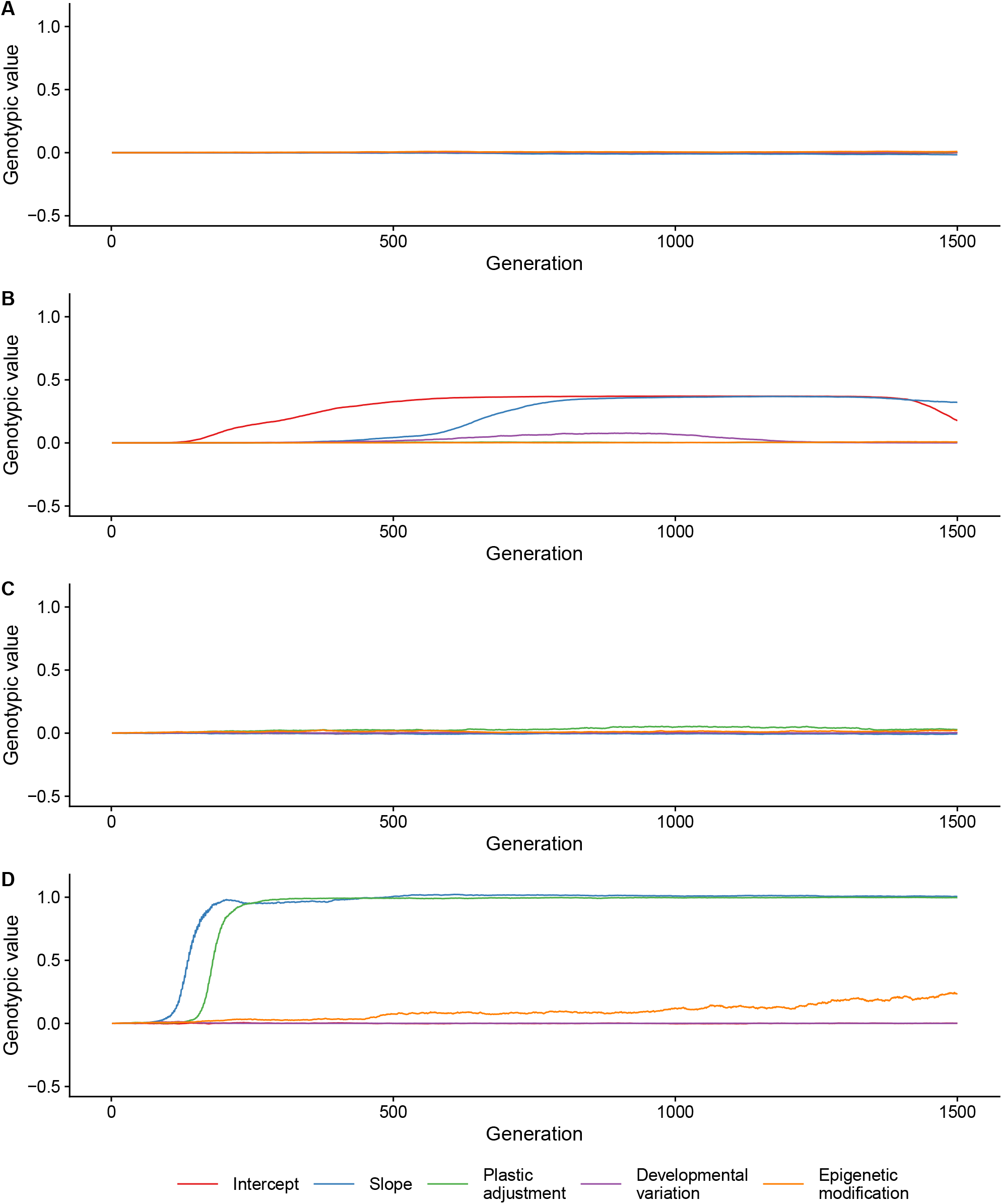
Examples of evolutionary trajectories when standard deviation of mutational effects is low. Genotypic values are shown for individual simulation runs. In all panels number of loci is 10 for each category, *P* = 1, *k*_*d*_ = *k*_*e*_ = 0.02, and *k*_*a*_ = 0.01. A) *R* = 1 (log10 *R* = 0), *σ*_*α*_ = 0.01 B) *R* = 3162.3 (log10 *R* = 3.5), *σ*_*α*_ = 0.01 C) *R* = 1 (log10 *R* = 0), *σ*_*α*_ = 0.1 D) *R* = 3.2 (log10 *R* = 0.5), *σ*_*α*_ = 0.1

